# An Empirical Bayes Method for Differential Expression Analysis of Single Cells with Deep Generative Models

**DOI:** 10.1101/2022.05.27.493625

**Authors:** Pierre Boyeau, Jeffrey Regier, Adam Gayoso, Michael I. Jordan, Romain Lopez, Nir Yosef

## Abstract

Detecting differentially expressed genes is important for characterizing subpopulations of cells. In scRNA-seq data, however, nuisance variation due to technical factors like sequencing depth and RNA capture efficiency obscures the underlying biological signal. Deep generative models have been extensively applied to scRNA-seq data, with a special focus on embedding cells into a low-dimensional latent space and correcting for batch effects. However, little attention has been given to the problem of utilizing the uncertainty from the deep generative model for differential expression. Furthermore, the existing approaches do not allow controlling for the effect size or the false discovery rate. Here, we present lvm-DE, a generic Bayesian approach for performing differential expression from using a fitted deep generative model, while controlling the false discovery rate. We apply the lvm-DE framework to scVI and scSphere, two deep generative models. The resulting approaches outperform the state-of-the-art methods at estimating the log fold change in gene expression levels, as well as detecting differentially expressed genes between subpopulations of cells.

## 1 Introduction

Single-cell omics measurements promise to resolve the heterogeneity of cellular identities, characterizing subtle molecular changes between cells [1]. In particular, measuring gene expression in single cells—with single-cell RNA-sequencing (scRNA-seq)—opens up the opportunity to characterize differences in complex scenarios, e.g., for rare cell-type detection or inter-cell-types/inter-conditions comparative analyses. However, technical factors such as the sequencing depth and batch effects obscure the underlying biological signal, requiring careful probabilistic modeling of the data to allow accurate detection of differentially expressed genes.

Many differential expression (DE) methods have been developed to handle the specificity of scRNA-seq data [2, 3, 4], often building on earlier work on bulk RNA-seq [5, 6]. However, there is a gap between the way these methods are applied and the structural characteristics of modern datasets. First, the performance of these approaches in modern datasets is highly dependent on their ability to correct for batch effects [7]. Indeed, a large number of these approaches rely on generalized linear models (GLMs), while recent benchmarking studies in scRNA-seq data integration focus on nonlinear methods (e.g., MNNs [8]) for their evaluation [9]. The linear method present in that benchmarking study, Combat [10] was found to be suitable only for simple scenarios. Second, most methods for DE analysis are unable to take advantage of the large sample sizes of current scRNA-seq experiments in learning their underlying gene expression model. Indeed, DE methods are often run on pairs of cell groups, subsampled from the original data. Gene expression level estimates could be improved by leveraging shared information across the whole dataset for model fitting, e.g., to refine a given gene expression levels from gene-gene correlation structures accross batches.

Recent advances in deep generative modeling [11] led to the development of a range of methods for normalization and analysis of gene expression data at scale, able to deal with millions of cells and able to account for complex study designs [12]. Most of these models (e.g., scSphere [13], scVI [14], scANVI [15], scGen [16] and CellBender [17]) represent the expected gene expression in cell *n* as a nonlinear function of a latent low-dimensional summary of the cell’s state *z*_*n*_ and its batch-identifier *s*_*n*_. Such approaches intrinsically lead to information sharing across cells and across genes and promise to address the aforementioned issues. However, little attention has been given to carrying out rigorous and accurate DE analyses with those models. Two exceptions are scVI [14] and scANVI [15] in which the uncertainty in the latent variable *z*_*n*_ is used to approximate a Bayes factor and detect differential expression. However, the Bayesian approach mentioned in [14, 15] has three fundamental limitations. First, the underlying definition of DE genes is too sensitive and may tag genes with negligible differences in expression and often of limited biological relevance. Second, interpreting Bayes factors is challenging, because of the difficulty to translate them to established multiple hypothesis testing metrics, e.g., False Discovery Rate (FDR). Third, scVI proposes to average Bayes factors from randomly sampled pairs of cells from each group, which lacks theoretical justification.

After presenting background information on differential expression and deep generative models (Section 2), we introduce lvm-DE, a general Bayesian framework for detecting differential expression derived from first principles, and that addresses those issues (Section 3). lvm-DE takes as input a fitted deep generative model of scRNA-seq data, a pair of cell groups and a target *α*. lvm-DE provide as output estimates of the log fold change for every gene, as well as a list of DE genes, with FDR control at level *α*. Notably, our Bayesian hypothesis formulation of differential expression uses a composite alternative, built from the log fold change (as in DESeq2 [5]) to avoid detecting spurious or lowly expressed genes. Then, we benchmark lvm-DE applied to scVI [14] and scSphere [13] against classical differential expression approaches on simulated and real-world datasets (Section 4). These results demonstrate that lvm-DE is a valuable tool for understanding differential expression effects in scRNA. An open-source implementation of lvm-DE is available in beta as part of the scvi-tools Python package [18] (https://github.com/scverse/scvi-tools/tree/pierre/DE).

## 2 Background

Before introducing our statistical framework, we describe two lines of work in scRNA-seq data analysis on which our work builds: differential expression and deep generative modeling.

### 2.1 Differential expression

Differential expression analysis of transcriptomics data aims to determine which genes have significantly different expression levels between two groups of cells *A* and *B* (e.g., representing different cell-types or cells obtained under different experimental conditions). It is known that the direct approach of comparing counts across groups is subject to noise and bias because of the various sources of technical factors that confound the data [19] (i.e., variation in library size, batch effects). To address this, all DE methods define (either implicitly or explicitly) a model with a latent variable 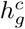 that represents the *denoised* or *normalized* expression level for each gene *g* and each condition *c* ∈ { *A, B* }. Differential expression for gene *g* between condition *A* and *B* is then assessed by comparing some statistic measuring the differences between the distributions of 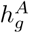 and 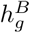. Finally, the significance of these differences is corrected to account for the multiplicity of hypotheses (one per gene) by estimating the false discovery rate (FDR).

Several popular DE frameworks for scRNA-seq data were originally developed for bulk RNA-seq data. Those methods are still competitive although not explicitly designed for scRNA-seq [20]. Two of the most prevalent methods, edgeR [6] and DESeq2 [5] model gene expressions as negative binomial distributions. Another widely used method, Limma-voom [21] uses log-transform counts instead. A common theme to all three methods, however, is that they rely on Generalized Linear Models (GLM), making it convenient to represent and linearly control for possible biases (e.g., due to batch effects or GC content). Although these GLMs are fit for every gene separately, it is important to note that their training procedures pool information across genes, relying on mean-variance relationships to improve the fit of some of the parameters using Empirical Bayes. Another key contribution, featured in DESeq2, is to propose a *composite* null hypothesis in which the hypotheses are made on the fold change being higher than some threshold. This results in exclusion of genes whose log fold change (LFC) is low, even if the differences in their expression are statistically significant. In practice, this procedure helps exclude genes with little biological interest.

Another set of DE analysis frameworks have been designed specifically for scRNA-seq data, with popular methods including MAST [2] and SCDE [3]. MAST uses a hurdle model that was designed to account for the abundance of so called dropouts in scRNA-seq data, namely zero counts that are observed mostly with lowly expressed genes, due to limitation in RNA capture efficiencies. SCDE takes a conceptually similar approach and employs a Bayesian framework to models scRNA-seq data as a mixture of negative binomial and Poisson distributions. The negative binomial component of the mixture accounts for counts that are attributable to gene expression while the Poisson component models dropout events [22]. In the following, we compare our approach for DGM-based DE against this collection of DE methods, coming from both bulk (DESeq2, edgeR, limma-voom) and the scRNA-seq literature (MAST).

### 2.2 Deep generative models for scRNA-seq analysis

With steadily growing sizes of datasets [23], recent work started to adopt more flexible and non-linear approaches for modeling gene expression in single cells. While these methods require more data for training, they are often complemented by scalable inference procedures. One popular line of work relies on advances in deep generative modeling [12], in which models make use of deep neural networks as link functions in their conditional distributions (e.g., [13, 14, 15, 16, 17, 24, 25, 26, 27, 28, 29]).

Most of these methods fit a hierarchical probabilistic model (i.e., a graphical model), that includes a low-dimensional latent variable embedding each cell in some cell state space.

To ensure that these cell embeddings reflect biological variation as much as possible, and remain disentangled from technical variation, a common practice is to model nuisance factors as parameters or random variables within the generative model. One factor that is commonly accounted for is variation between cells due to sequencing depth. For instance, scVI [14] and scANVI [15] models include a random variable that represents the sequencing depth of each cell. This variable is used as a scaling factor, thus providing normalized expression levels. The hierarchical model also links the latent representation of cell state directly to the normalized expression values, thus decoupling the cell embeddings from variations in sequencing depth. An alternative approach, scPhere [13] uses the sum of observed transcripts as a deterministic scaling factor, but relies on an additional penalty term to ensure that the cell embeddings are decoupled from it. Deep generative models provide accommodating strategies to correct for other nuisance factor(s) present in single-cells. Multiple approaches, including scVI, scANVI, or scPhere assume model normalized gene expression as a function of both cell state embeddings and these nuisance factors. This function learned during the fitting process and parameterized as a multi-layered perceptron can model intricate non-linear nuisance effects on gene expression. Such nuisance factors can include batch or patient id in multi-donor data sets, but have also been generalized to broader covariates, e.g., cell cycles [18].

Distributional assumptions are other key components that can improve cell-state representation and gene expression modeling. For example, in scGen [16], the counts are normalized, log-transformed and treated as samples from a normal distribution. Conversely, both scVI [14] and ScPhere [13] model gene expression counts under a negative binomial likelihood. Other models (e.g., the original work of scVI) include the addition of an additional low-magnitude term, supposedly handling zero-inflation in the data, due to limitations in sensitivity and the effects of transcription bursting (discussed in more depth at [30,31]). Another component of importance is the modeling of the cell embedding. DCA [26] relies on a deterministic embedding, treated as a parameter optimized using maximum likelihood estimation. Another line of work, variational autoencoders (VAEs) posit a prior distribution on the embedding that regularizes the embedding space and aims to learn more useful representations [11]. While most unsupervised VAEs posit a non-informative isotropic Gaussian prior, alternative priors may improve cell representation in specific contexts. For instance, scANVI [15] leverages partial data annotations to learn more structured priors on the latent variable. scPhere embeds biological states in a hyper-spherical space with a uniform prior, which could improve the modeling of rare cell types and hierarchical structures.

While most applications of deep generative models in single-cells revolve around the cell state space, few approaches consider normalized gene expression values and their uncertainty, that are central for DE. Aiming in this direction, DCA feeds denoised gene expression predicted by the model as inputs for DESeq2 instead of raw counts to better reduce the noise present in scRNA-seq data sets. This is unsuitable from the computational perspective because DESeq2 may be too long to run when dealing with groups of tens of thousands of cells, creating a bottleneck. Also, as DCA relies on an autoencoder, it does not provide uncertainty estimates on normalized gene expression, and summarizes the data to its mean instead. This is known to potentially create spurious signal in other downstream applications (e.g., gene-gene correlation estimation [32]).

scVI and scANVI (e.g., [14, 15]) both provide integrated tools to perform differential expression between two groups. First, expected expressions are sampled from the posteriors of representative cells. These values are then subject to Bayes factor computation to assess significance. This framework has several limitations. First, although Bayes factors are an appealing alternative to frequentist hypothesis testing for Bayesian models, they may be hard to interpret while analyzing real-world data and cannot be employed towards decisions with false discovery rate control guarantees. Second, these approaches estimate significantly DE genes without considering the effect size (i.e., LFC), which can be extremely low for lowly expressed genes, relatively abundant in scRNA-seq data. Third, this approach is convenient since model fitting is performed only once using all cells, thus avoiding the need to refit a new model for each DE test (each considering different pairs of cell subsets). However, scVI does not explicitly account for cell type in its generative model. As a heuristic, scVI aggregates cells by averaging approximate Bayes factors across randomly sampled pairs of cells to compare cell populations. We provide more theoretical insight, and an improved approach in this manuscript.

## 3 Methods

We introduce lvm-DE, a principled framework to enable the use of deep generative models as means for quantifying and assessing the significance of differences in gene expression (Figure 1A). lvm-DE takes as input a latent variable model, a scRNA-seq dataset with covariates (such as batch identifiers) and two groups of cells to be compared, as well as a significance level *α*. lvm-DE provides an estimate of the LFC of each gene, between cells of the two groups. It also provides a list of significantly differentially expressed genes that controls the FDR at level *α*. These estimates are derived while controlling for differences in library sizes and for variability due to observed technical confounders, such as the association of samples with different batch identifiers.

**Figure 1:**
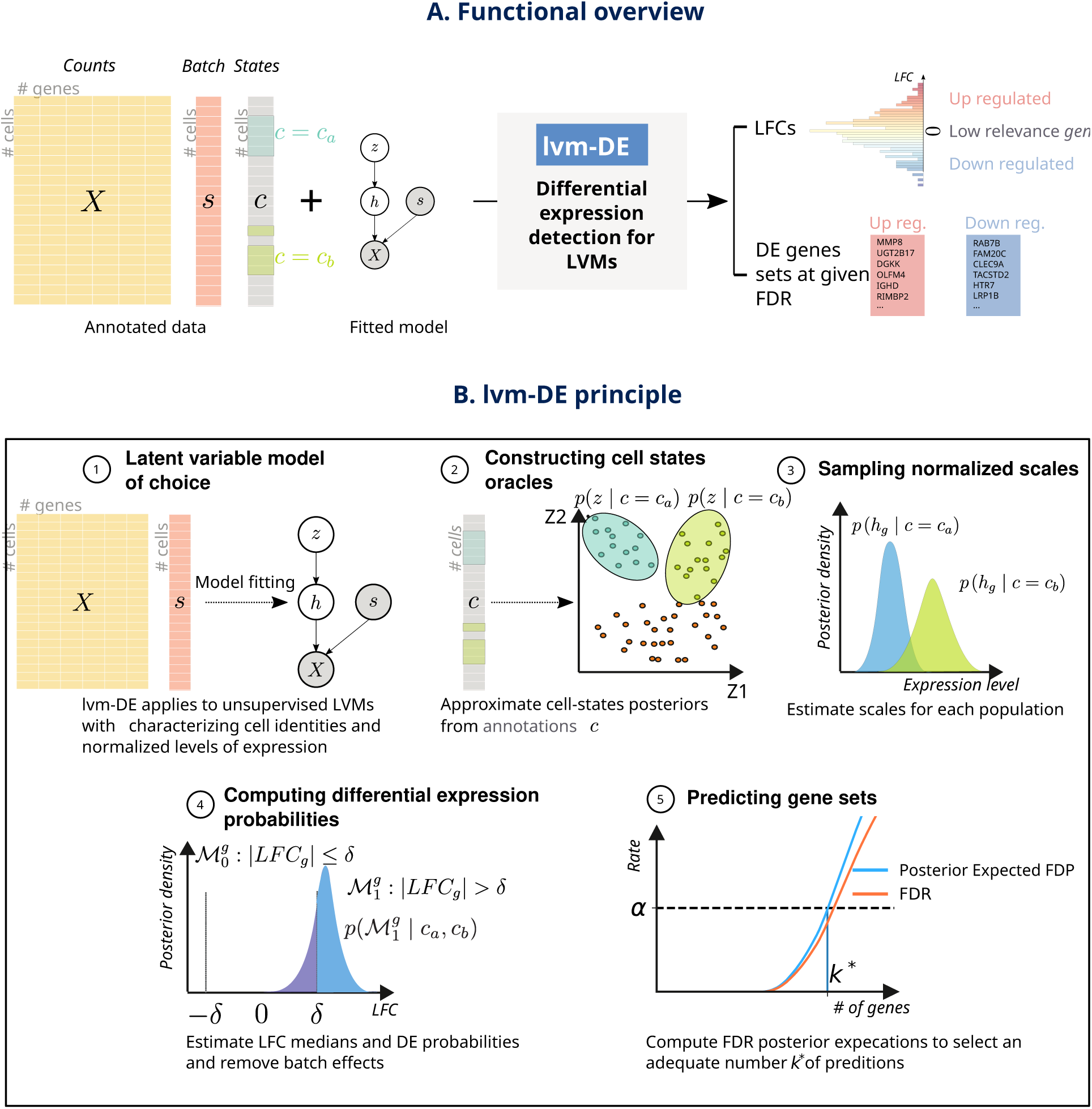
Differential expression model for deep generative models. **A**. lvm-DE takes annotated data (from clustering, metadata, or transfer learning), a latent variable model, and a target FDR level as inputs and returns log fold change (LFC) estimates as well as calibrated DE predictions. **B**. lvm-DE works as follows. 1. A preliminary step consists in fitting the latent variable model of choice of the data from the collection of available scRNA-seq data. 2. lvm-DE uses existing cell states annotations to approximate the distributions of *c* conditioned on the cell-states. 3. These distributions help determine the normalized expression level distributions of the compared populations. 4. The associated LFC distribution helps to determine posterior DE probabilities that correspond to the model in which the LFC is higher than a given threshold. 5. To tag DE genes of interpretable interest, we estimate the maximum number of genes for which the posterior expected FDR is below the desired FDR level.

lvm-DE requires five steps (Figure 1B). First, a model is fitted, such that the observed expression values in each cell (X) are represented by a low dimensional latent random variable (often denoted as *z*; in VAE-based models such as scVI, this amounts to learning an encoder network). This representation provides a summary of the state of each cell, while controlling for differences due to batch effects (s) and sequencing depth, but it does not use the annotation of cells into the two groups that we wish to compare (Figure 1B1, Section 3.1). Performing DE with an annotation-agnostic model is convenient as it does not require refitting the model when the we want to compare different groups of cells. Second, annotations are used to build a posterior distribution of the latent representation of each of the two groups, providing a summary of their characteristic cell states (Figure 1B2), Section 3.2). Third, cell states posteriors are used to infer a distribution of normalized expression for every gene (Figure 1B3) in each cell group. These high dimensional distributions are generated by propagating the low-dimensional representations through the latent variable models (LVM). In VAE-based models, this amounts to applying the decoder network. Fourth, the normalized expression values are used to estimate the LFC and a posterior probability for the LFC to be larger than a given cutoff *δ* (Figure 1B4, Section 3.3) while correcting for batch effects (Section 3.4). Finally, these posterior probabilities of DE are used to call genes whose significance is within a desired FDR *α* (Figure 1B5, Section 3.5).

### 3.1 Selection of deep generative models for lvm-DE

Our formulation of lvm-DE is general, such that it applies broadly to many types of deep generative models, and can operate as a wrapper function for differential expression with these models. In the following, we provide the modeling prerequisites for using lvm-DE and then discuss additional considerations of modeling choices that can lead to better accuracy.

Let *x*_*ng*_ be the observed numbers of transcripts in cell *n* that originate from each gene *g* and let *s*_*n*_ correspond to the batch annotation. A general form of generative process that qualifies for use by lvm-DE is as follows. First, a low-dimensional latent variable *z*_*n*_ representing a cell’s biological state is sampled from a prior distribution. A neural network then takes a sample from *z*_*n*_ and the batch information *s*_*n*_ and maps it to another latent variable, *h*_*ng*_ representing the underlying (i.e., normalized) expression level for gene *g*. Finally, the observations *x*_*ng*_ are drawn from model-specific conditional likelihood distributions, such as negative binomial or log-normal, with or without zero inflation [14, 16, 33]. The likelihood of a single observation (cell) in these models should decompose as follows:

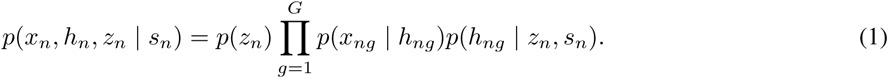

Furthermore, lvm-DE assumes that there exists a tractable approximation to the posterior *p*(*z*_*n*_ | *x*_*n*_) (e.g., using variational approximations [11], or MCMC [34]). While many published methods satisfy this basic prerequisite, here we selected two such models: scVI and scPhere [13, 14], both of which are based on a VAE but differ in their loss function and distributional assumptions.

Beyond this basic prerequisite, we identified two considerations for improved performance (implementation details in Appendix A). The first consideration is the need to model the normalized expression levels *h*_*ng*_, while controlling for nuisance variation in sequencing depth. In scVI and scPhere, this is addressed by modeling the gene expression count *x*_*ng*_ as a negative binomial distribution with mean parameter *l*_*n*_*h*_*ng*_, where *l*_*n*_ is a normalizing factor that can be interpreted as the inferred library size. The second consideration stems from the fact that lvm-DE exploits the variational distribution (i.e., an approximation of *p*(*z*_*n*_ | *x*_*n*_)) for Bayesian hypothesis testing. In this setting, it has been reported that VAE trained with the classical evidence lower bound (ELBO) may underestimate the variance of the variational posterior and thus under perform in tasks of decision making, compared to models trained with alternate variational bounds [35]. Consequently, we train scVI and scPhere using the importance-weighted evidence lower bound (IWELBO), which was demonstrated to lead to improved performance in other application domains [36].

### 3.2 Aggregating individual cell information to capture population expression levels

lvm-DE takes as input a stratification of cells into groups (*c*_*n*_ ∈ { 1, …, *K* }; e.g., representing different cell types or different conditions under which cells were captured), along with a generative model that conforms with Equation 1.

To estimate the distribution of gene expression in every group of cells, we assume that there exists an oracle function Φ mapping a point in latent space *z* to the corresponding covariate *c* = Φ(*z*) ∈{ 1, …, *K* }. This is a reasonable assumption because the low-dimensional representation given by scVI or other deep generative models often successfully stratifies cells according to their groups [14].

Given a particular value *C* of the covariate *c* (e.g., a cell type or a condition), we denote by 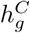 the normalized expression level of gene *g* in condition *C*. We define 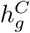, conditioned on a given batch *s* as

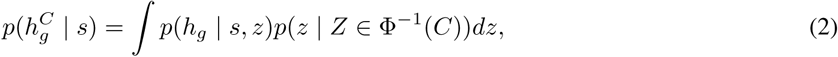

where *p*(*z* | *Z* ∈ Φ^*−*1^(*C*)) denotes the prior density *p*(*z*) truncated to the preimage of { *C* } under the oracle function Φ. Because Φ is unaccessible to us, we may estimate the prior density from the data, in the spirit of empirical Bayes methods. More precisely, we first approximate it using the posterior of the annotated data:

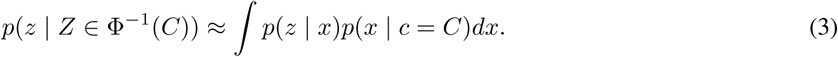

Second, we estimate this quantity using the truncated dataset with cells belonging to condition *C*, yielding the mixture density

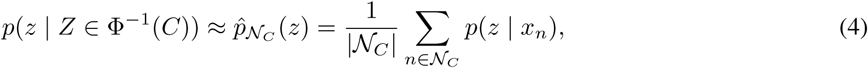

where 𝒩_*C*_ is the set of cell indices that belong to the condition *C*. To reduce the possible sensitivity to outlier cells, we developed a procedure to identify and remove these from the set 𝒩_*C*_. To do so, assuming that the latent mean representations within a cluster are normally distributed, we estimate the covariance matrix from posterior mean samples 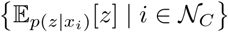, and remove observations whose variational mean falls outside the 90%-confidence ellipse described by the covariance estimate. As demonstrated previously, the empirical prior appearing in Equation 4 is intractable. A first solution is to replace the individual posterior distributions *p*(*z* | *x*_*n*_) with their variational approximations *q*_*ϕ*_(*z* | *x*_*n*_), providing the so-called plugin estimator

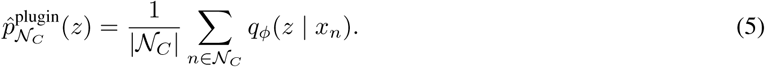

A remaining caveat of this estimator is that it lends equal importance to each sample from the cell population and of the variational distribution. This is in a way similar to the original scVI, in which pairs of cells are sampled uniformly at random from the two populations and compared to each other. It is reasonable to expect, however, that in practice some samples may be of significantly better quality, which explains the previously reported increased performance from using importance-sampling based estimators [35]. Based on this observation, we use the mixture prior from Equation 4, approximated by self-normalized importance sampling (SNIS) with the mixture of Equation 5 as the proposal distribution. In this formulation, we first sample 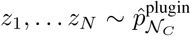, and the contribution of each sample *z*_*i*_ will be proportional to:

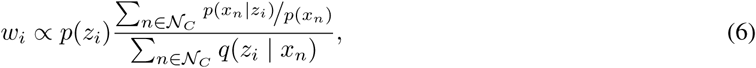

where *p*(*z*_*i*_) denote the prior density for *z*_*i*_, and the intractable evidence terms *p*(*x*_*n*_) are estimated with IWELBO estimators (*K* = 5, 000 particles). The final samples 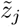 correspond to random samples with replacement from {*z*_*i*_, *i* ≤ *N* }, where each entry *i* of the set has weight 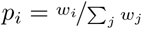 [36], such that

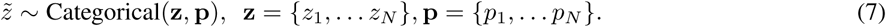

Consequently, samples from *z* that have high likelihood levels across a large pool of cells from 𝒩_*C*_ will have higher weights in the posterior, thus further improving the robustness to outliers and to misassignment of cells into groups. In order to ensure that the importance sampling procedure is sufficiently stable, several diagnosis tools such as the Pareto smoothed importance sampling (PSIS) estimator [37] may be employed. Once we have samples from the distribution *p*(*z* | *Z* ∈ Φ^*−*1^(*C*)), we may follow Equation 2 to link these samples to the normalized expression levels 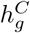 for DE analysis. In the case of scVI and scPhere, expression levels *h* and cell states *z* are linked by a neural network *h* = *f* (*z, s*).

### 3.3 Fold change estimation for DE

A practical limitation of DE analysis is the identification of genes in which the differences in expression are significant and yet of low magnitude (low LFC), and thus of less or no interest. To address this, lvm-DE treats the LFC itself as a random variable and uses a Bayesian analysis to retain only genes in which the effect size is significantly larger than a cutoff.

To quantify our confidence that the LFC is beyond a desirable level, we introduce the effect size random variable,

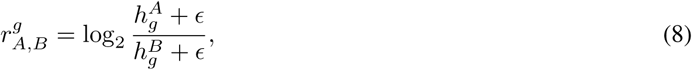

where *ϵ* is a small offset used for stability. When gene *g* has very low expression levels in both conditions, the pseudo-count *ϵ* ensures that the associated LFC has a low magnitude. More details about the choice of *E* can be found in Appendix C).

To test for differential expression in lvm-DE, we introduce three models for which gene *g* is either up-regulated 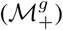, down-regulated 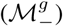 or not differentially expressed 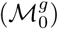 in population *A* compared to *B*. These three models are defined via the change variable 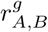 as follows:

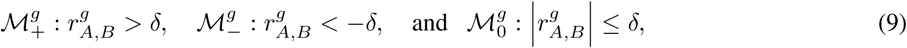

where *δ* is a threshold ensuring that the selected genes are interesting in practice. We discuss heuristics for the choice of *δ* in Appendix D). The probability of each of those models may be estimated from the data, using the empirical Bayes method and the SNIS estimator. In particular, we note:

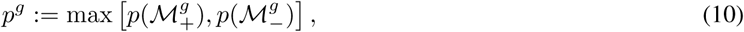

the probability that gene *g* is DE (i.e., with absolute LFC greater than *δ*).

The LFC is a widely used measure of effect size for expression measurements because of its interpretability [5, 38, 39]. However, our framework could be extended to any difference measure between expression levels, including logit fold change for DNA methylation, as in scMET [40].

### 3.4 Batch effect correction

Our exposition in the previous section assumed that all the cells come from a single batch. However, DE analyses are often required in multi-batch or multi-donor settings.

Two scenarios appear when performing differential expression, depending on whether all batches contain cells from both conditions. In the most natural setting, such as a biological replicates, we expect a representation of all the compared conditions in all batches. In this case, let *S* be any set of batch indices for which there exists at least one observation of each condition. Following our Empirical Bayes approach, we define batch-specific empirical priors relying on the subset of observations 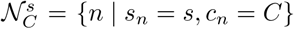. Then, the probability of each of the three models may be computed by averaging across batches (similar calculations were proposed in [15]). For example, the probability of up-regulation is computed as

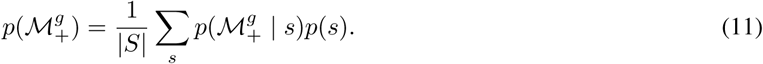

In some setups however, there may not be perfect overlap of each condition in all batches (e.g., when comparing tumors, or inflamed tissues to healthy controls). In this more ambiguous scenario, we condition the posterior scales distributions w.r.t. observed batches, i.e., 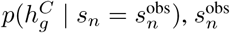 being the batch that cell *n* originates from.

### 3.5 Controlling the false discovery rate

Bayesian decision theory [41] shows that the optimal decision rule for our DE task consists of selecting the top genes ranked by *p*^*g*^ (Equation 10). However, selecting the number of “top” genes is not straightforward. The use of Bayes factor to guide this choice carries the complication of not being interpretable when it comes to the expected error rates. A more intuitive solution is therefore to set the significance threshold *α* over *p*^*g*^ in a way that controls for the false discovery rate (FDR).

To this end, let *d*_*g*_ be the (unknown) binary random variable denoting whether or not gene *g* is DE. Consider the multiple binary decision rule 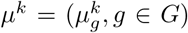 that consists in calling DE the *k* genes with the highest differential expression probability *p*^*g*^. The false discovery proportion of the decision rule *μ*^*k*^ can be expressed as

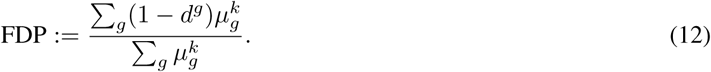

We use the formulation of the posterior expected FDP, as introduced in [42] and applied to genomics data in [43, 44], corresponding to

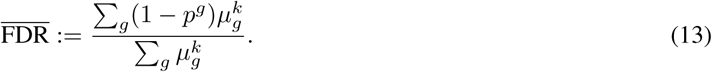

Because this quantity is tractable after inference, a natural choice for *k* corresponds to the maximum value for which the posterior expected FDP is below the desired level *α*.

## 4 Results

lvm-DE addresses two key tasks in differential expression analysis. The first corresponds to the robust estimation of the LFC in noisy and sparse data, including for rarely observed cell states.. The second task is evaluating the statistical significance of the observed effect size, and calling genes as differentially expressed for any target false discovery rate. For both tasks, we apply the lvm-DE framework to scVI and scPhere. We denote scVI-lvm and scPhere-lvm the associated combinations, in contrast to scVI that natively provides a test for differential expression. We evaluate our models in comparison to the current state-of-the-art, including MAST [2], DESeq2 [5], edgeR [6], and Limmavoom [21] (referred to as voom). For each of these models, we follow the guidelines and implementation details specified in a recent comparative study of DE methods for scRNA-seq [20] (further details in Appendix E). We assess lvm-DE performance for LFC estimation, and for the prediction of correct and calibrated DE gene sets using both synthetic and real datasets. In the real datasets, we focused the analysis on the top 3,000 highly variable genes using Seurat. Annotations were taken from the respective publications.

### 4.1 Accurate and calibrated results on simulated scRNA-seq data

As a simulation scheme, we employ SymSim [45], which models biological and technical noise factors (e.g., transcriptional bursting and PCR amplification), as well as statistical dependencies between genes (relying on low dimensional representation of the kinetics of promoters in each cell). We used SymSim to generate a hierarchy of cell states and focus our analysis on the comparison of two “sibling” types, denoted A and B (Figure 2A). We generated a dataset of 10,000 cells and 1,000 genes across five cell-types. The data corresponds to two batches (5,000 cells each) with unique molecular identifiers (UMIs), which have similar properties to 10x Chromium data. To estimate the ground-truth LFC, we use the expected gene expression means, derived from the expected promoter-kinetic parameters in each gene and cell-type [45]. The resulting dataset contains a total number of 396 genes, including 127 differentially expressed genes. We selected these genes such that DE genes have non-ambiguous high expression differences (absolute LFC higher than 1), while negatives have negligible expression differences (absolute LFC lower than 0.2).

**Figure 2:**
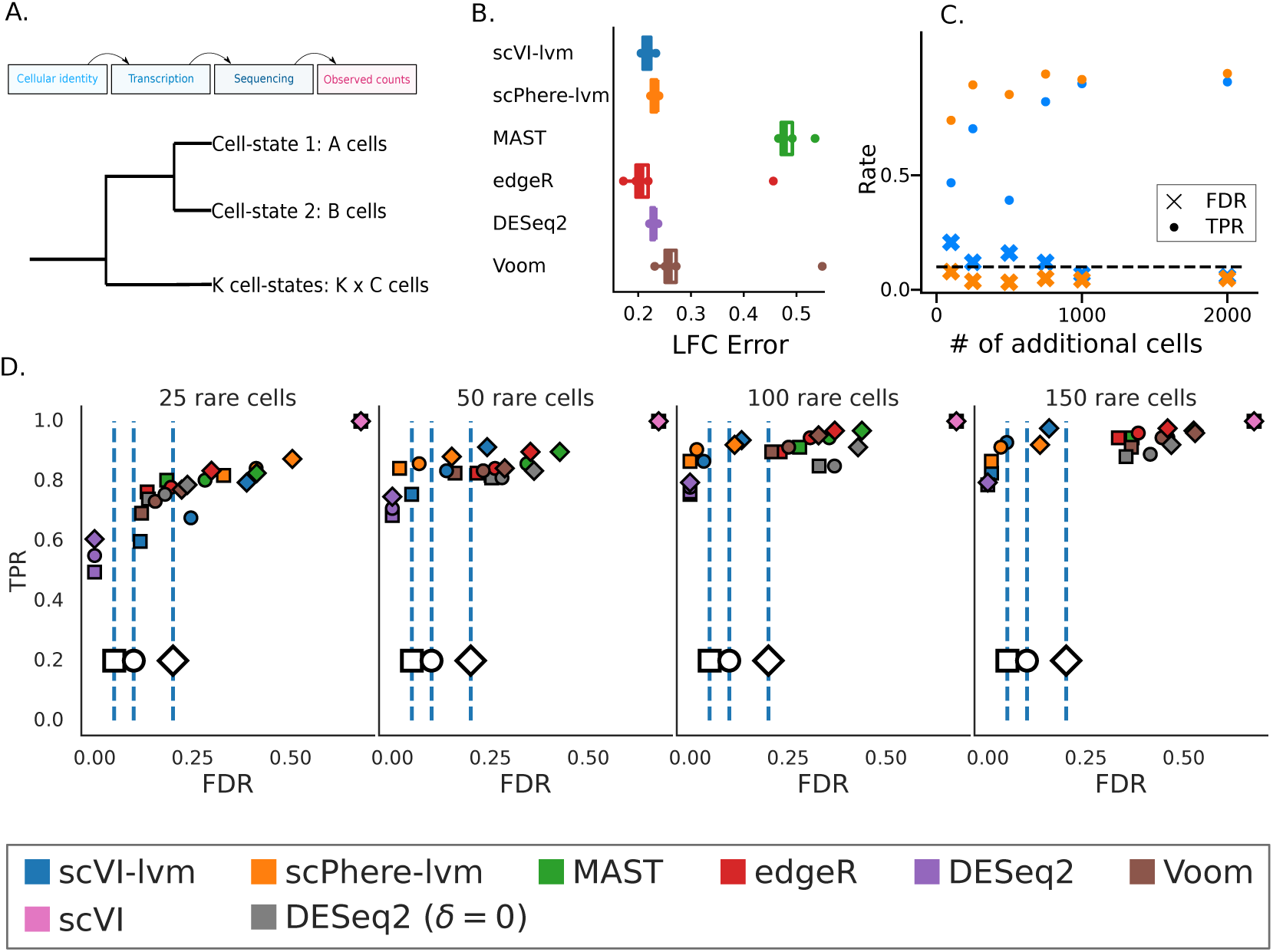
SymSim results. **A**. Dataset presentation. *Top*: SymSim is a simulation framework modeling biological and technical effects to provide realistic simulations. *Bottom*: We consider a two cell-type DE analysis scenario. We subsample population A to compare the different algorithms for rare cell-type detection. For this sub-panel and all the experiments, we refit the models for each scenario such that all of the algorithms use the same number of observations from A and B for model fitting and DE. **B**. LFC point estimation error when comparing two populations of *A* = *B* = 200 cells. For Bayesian techniques, we summarize the posterior LFC distribution by its median. For this figure and in the remainder of the article, boxplots represent medians (line), interquartile range (box), and distribution range (whiskers) estimates. **C**. TPR (*dots*) and FDR (*crosses*) changes for an increasing number of external cells for the different latent variable models. **D**. FDR and TPR of decisions for the detection of DE genes when comparing varying *A* ∈ {25, 50, 100, 150} cells to *B* = 500 cells (for *C* = 2000). Squares, circles, and diamonds correspond to decision controlling FDR at targets 0.05, 0.1, and 0.2. For scVI’s original DE procedure, we reject the null when Bayes factors are greater than 3 in absolute value.

We first evaluated LFC estimates from the models. The lvm-DE-based models offered accurate estimates (Figure 2B), on par with edgeR and DESeq2. This result suggests that lvm-DE properly aggregates individual cells to infer expression levels in the two subpopulations. An outlier in this analysis was MAST that exhibited a high mean-squared error range. This likely comes from the fact that this method relies on a hurdle model, whose LFC is difficult to compare to the ground-truth LFC. It was therefore discarded from further LFC estimation experiments.

By definition, scVI-lvm and scPhere-lvm make use of all cells in the input data, including “external” cells that are not in the groups being compared. In our simulation these are simulated cells that are not in groups A or B (Figure 2A). To test whether the inclusion of more external cells affect accuracy, we varied the number of external cells and conducted differential expression analysis between groups A and B (which remain unchanged). We find that our FDR control procedure is well calibrated, and remains at the expected level in all cases. Furthermore, we find a steady increase in the true positive rate (Figure 2C). These results indicate that increase in the overall data sets size, but not necessarily at the compared populations, leads to increase in power, thus supporting the notion the DGMs can benefit from information on all cell types (e.g., gene-gene correlations) in a dataset. This is in contrast with the other benchmark methods listed here which do not absorb any information from cells outside the compared groups.

The results in Figure 2D support the suitability of lvm-DE for characterizing rare cell-types, as capturing gene couplings can help alleviate the issue of small sample size. We tested this by leaving in the input data only a few cells of group *B* (representing a rare cell type), while keeping group A as before (representing a more prevalent background; Figure 2A). For a rare subpopulation of size 50 and up, both lvm-DE methods provide gene lists that are accurate and control the FDR for several significance levels. For the GLMs, we observed acceptable FDR control in low sample size setups, but interestingly their FDR becomes higher than its expectation, with larger sizes of the rare subpopulation. An exception is the default DESeq2 procedure, which uses a composite null hypothesis (i.e., effect size threshold *δ >* 0). This method does not suffer from large FDR, but loses power because of the difficulty to properly set the effect size threshold in practice. Finally, we observe that the Bayes factor approach originally used in scVI reaches high FDR levels. Consequently, we excluded this procedure from the next experiments.

To further our confidence in these results, we expanded the analysis in two ways. In the first analysis, we modified our simulation process to generate additional, and possibly more nuanced scenarios of differential expression, such as bimodality of the ground truth gene expression [46]. Our procedure compares favorably to the other algorithms as it provides a high number of true positive genes, and few false positives (Figure S1). In a second analysis, we used MUSCAT [47] as our simulation framework. Unlike SymSim, the data generated by MUSCAT is based on independent sampling from an underlying (here, negative binomial) distribution. Therefore, in this case, there is no benefit in using latent variable models, as no couplings between genes (which is to be expected in real data sets) can be leveraged. In such setups, the lvm-DE methods provided calibrated predictions, while having similar or lower level of power compared to the GLMs, as expected (Figure S2).

### 4.2 Favorable comparison to current methods on single-batch, medium-sized PBMC data

To test the performance of lvm-DE with data, we used a 10X Chromium dataset of 9,432 peripheral blood mononuclear cells (PBMC) from a human donor [48] (Figure 3A). This data provides an opportunity to establish the relative performance of lvm-DE in a configuration in which batch effects are not present.

**Figure 3:**
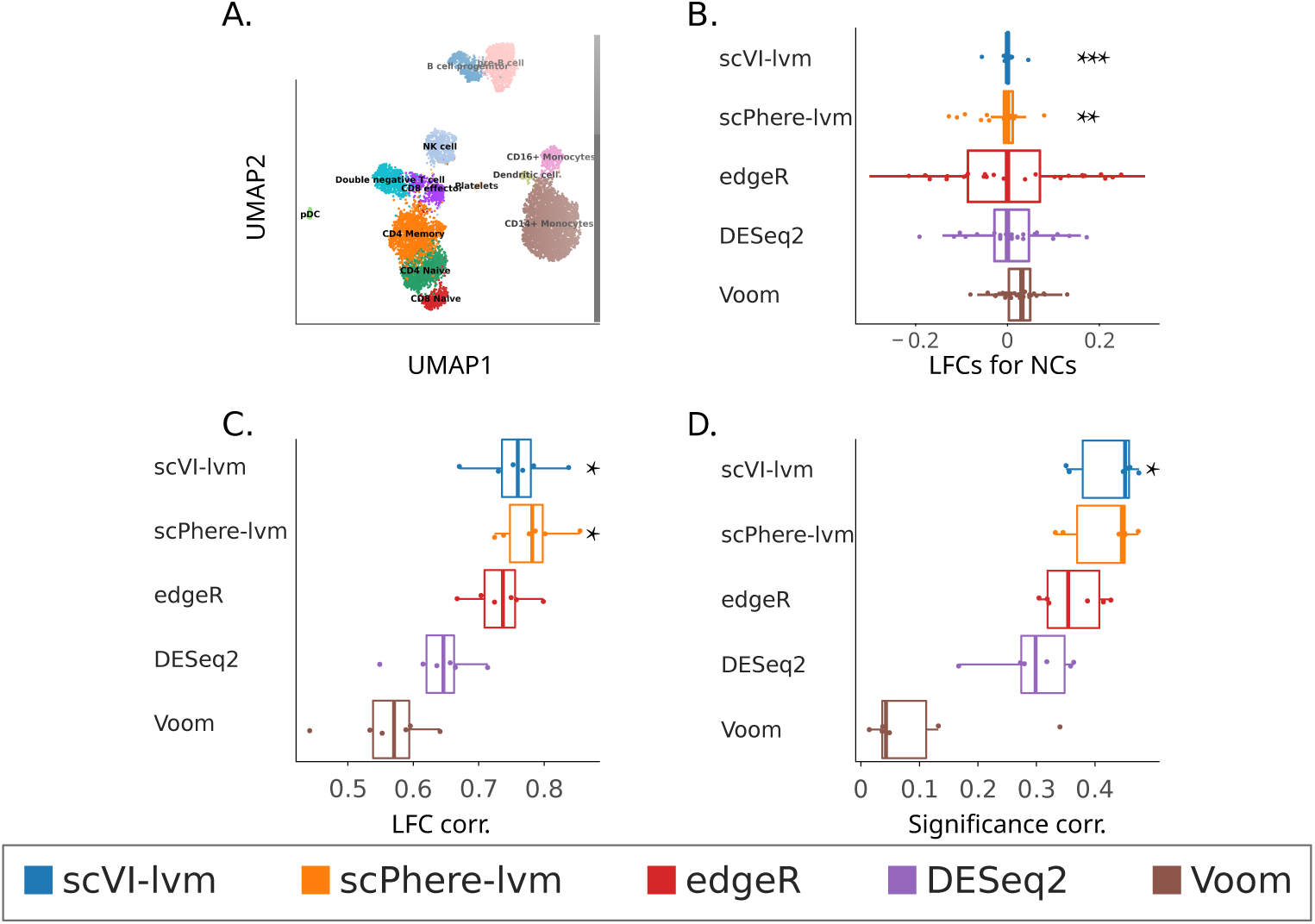
PBMCs results. **A**. UMAP from scVI’s embedding **B**. Negative controls (among B cells), corresponding to LFC range study for the different methods. For this experiment, the lvm-DE outlier removal procedure was not employed. **C**. Positive controls. Distribution of Pearson correlation between the reference LFC (bulk-RNA) and estimated LFC for pairwise comparisons of B cells, mDC, pDC and monocytes. Each point in these graphs corresponds to one of the six possible cell-type comparison. For lvm-DE, we use the custom LFC median estimator. Individual scatter plots can be found in the annex. **D**. Distribution of Spearman correlations between the reference pvalues (bulk-RNA) and estimated significance scores for pairwise comparisons of B cells, mDC, pDC and monocytes. GLMs and lvm-DE respectively used pvalues and posterior DE probabilities as significance scores. Stars represent significant differences with all the GLMs at various significant levels (*, **, and *** denote respectively significance levels *<* 0.05, 0.01, 0.005), under a two-sample F test for the negative control and a one-sided two-sample t test for the positive control experiments.

While there is no ground-truth for differential expression in this data, several strategies can help evaluate the estimation of differential expression. First, we can ensure that lvm-DE does not detect differences in expression when comparing groups of cells of the same state. We therefore construct negative controls (Figure 3B) by randomly splitting the B cell population (460 cells) into two groups (230 cells each) and predicting the LFCs with the different algorithms. In this configuration, we expect to measure negligible expression differences. lvm-DE LFC estimation compares favorably to its competitors, since scVI-lvm along with scPhere-lvm predict LFC of significantly lower amplitude than their GLMs counterparts.

Another benchmark strategy is to compare the outcome of the DE analysis to results obtained from an independent data set. To this end, we used estimations of LFCs and p-values from a bulk-RNA-seq dataset [49], in which DESeq2 was used for pairwise comparisons of monocytes, B cells, plasmacytoid, and myeloid dendritic cells.

The bulk-level LFCs and significance scores were then compared against the results of applying the different algorithms on the single cell data (Figures 3C and 3D). We find that scVI-lvm and scPhere-lvm compare favorably to the GLMs in recapitulating both the LFC and the significance scores of the reference bulk data. Notably, the cell groups we analyzed had varying sizes (Figure 3A), demonstrating that lvm-DE can capture state differences for abundant as well as less common cell-types. A detailed comparison of LFC estimations for every method and for every pair of cell types is depicted in Figure S3.

### 4.3 lvm-DE pools information from different sequencing technologies for better differential expression

lvm-DE can be employed to perform DE in large-scale multi-batched data. As an illustration, we consider the PbmcBench data collection [50], consisting of two biological samples sequenced with seven different protocols, either droplet (drop-seq, 10Xv2, 10Xv3, inDrop and CEL-seq) or well-based (seq-well and Smart-Seq2). This data provides the opportunity to further validate the ability of our procedure to alleviate technical factors’ impact on DE.

We compare the estimated LFC of scVI-lvm with bulk-RNA using the same reference as in the previous experiment. The inclusion of additional data sequenced with different technologies consistently increase the correlation between lvm-DE effect-size and significance predictions with bulk-RNA data (Figure 4). In contrast, observing more replicates does not systematically improve the match of GLM’s predictions with bulk-RNA. This suggests that contrary to GLMs, lvm-DE succeeds in pooling information from heterogeneous experiments to better capture differences between cell-states.

**Figure 4:**
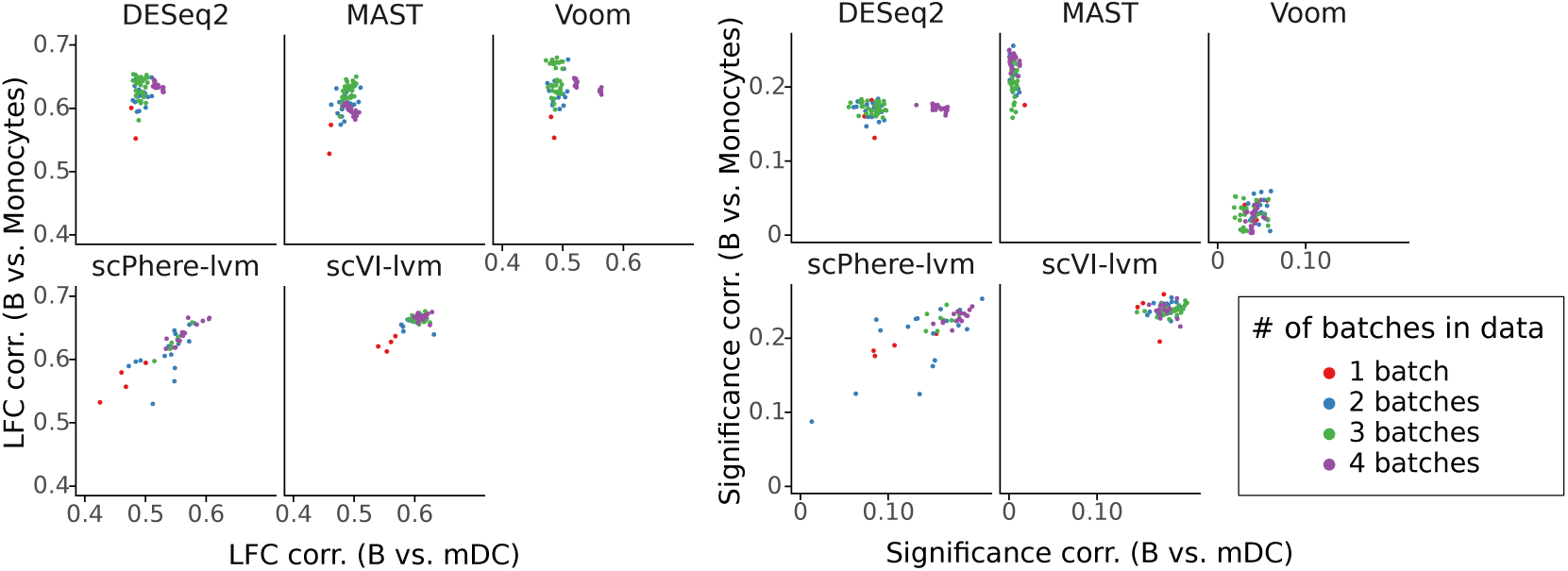
Batch harmonization on *PbmcBench*. Pooled information from several batches improves the match of the prediction with bulk for scPhere and scVI-lvm. *Left*: Pearson correlation of the predicted and the reference LFCs (from bulk-RNA) for two cell-type pairs. *Right*: Spearman correlation of the predicted significance scores (posterior differential expression probabilities for scPhere and scVI-lvm, pvalues for other algorithms) and the reference pvalues (from bulk-RNA) for two cell-type pairs. In both graphs, points correspond to a given training on a subset of *PbmcBench* containing a varying number of batches (color). As GLMs struggled to scale to large sample sizes, these algorithms used a maximum of 500 cells per dataset.

In summary, lvm-DE benefits when either the number of observed cells increases or when more samples are available, even if the samples differ technically.

### 4.4 lvm-DE better recapitulates inflammatory gene expression patterns in COVID-19 patient data

We now focus on a larger PBMC dataset, consisting of 44,721 cells from seven healthy donors and six patients with confirmed SARS-CoV2 infection [51] (Blish). This dataset provides an opportunity to evaluate procedures for differential expression, not only at comparing different cell types (to find their respective markers), but also comparing donor groups *within* each cell-type. The latter comparison, revealing the influence of inflammation on different immune compartments, is usually more challenging with a more nuanced and less reproducible signal [52] (Figure 5A).

**Figure 5:**
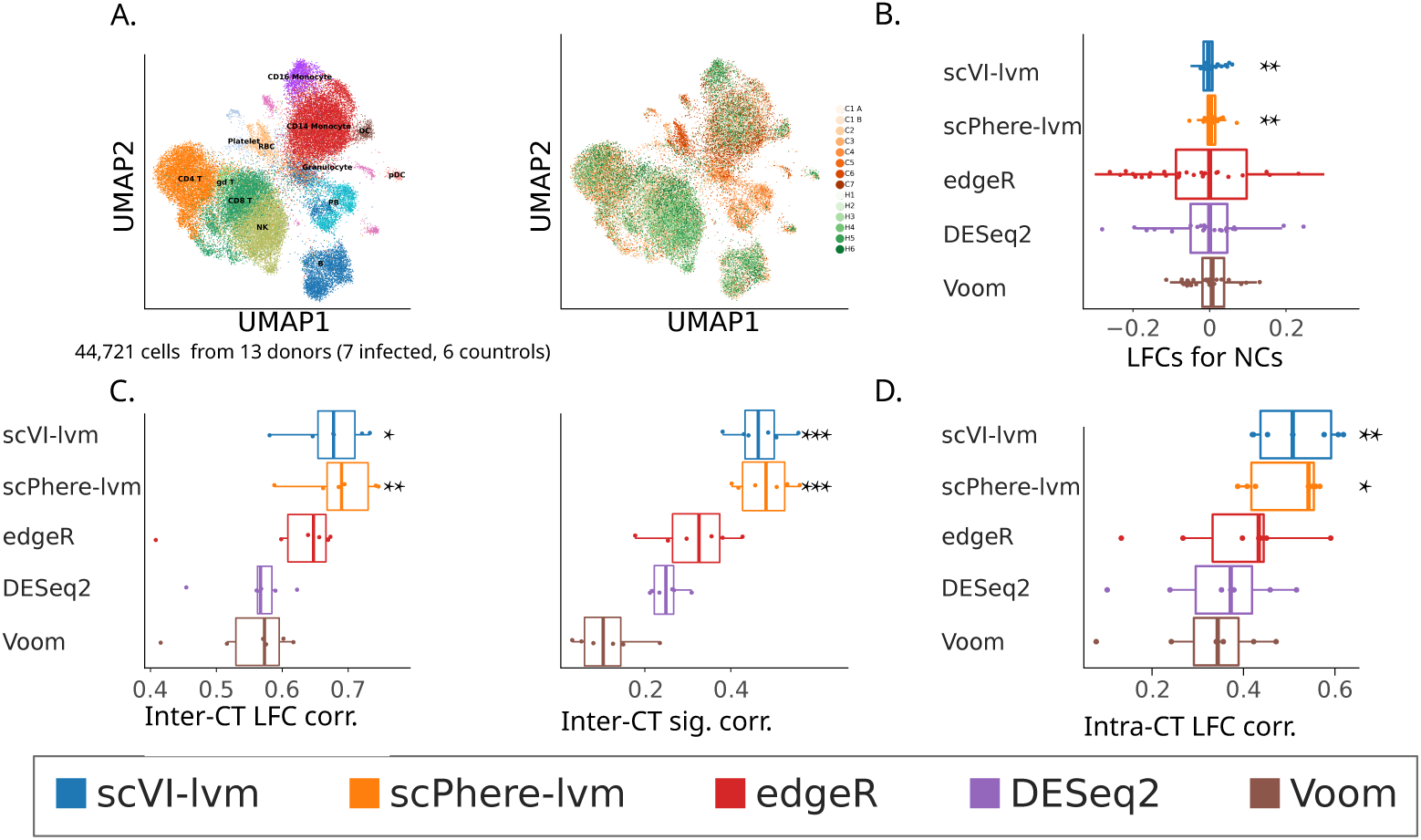
SARS-CoV2 dataset results. **A**. Dataset presentation. UMAP from scVI’s embeddings colored by cell-type (*left*) and batch (*right*). Counts were obtained from 6 healthy donors (H1 to H6) and 7 SARS-CoV-2 infected patients (C1 to C7). **B**. Negative controls (among DC cells), corresponding the study of the range of the LFC parameter (LFC) for the different methods.. **C**. Positive controls for inter cell-type analysis. *Left*: Distribution of Pearson correlations between the reference and estimated LFC for pairwise comparisons of B cells, mDC, pDC and monocytes. *Right*: Distribution of Spearman correlations between the reference pvalues and estimated significance scores for pairwise comparisons. Each point in these graphs corresponds to one of the six possible cell-type comparison. **D**. Positive controls for within cell-type analysis. Distribution for different cell-types of Pearson correlations between the reference and estimated LFC. The reference corresponds to cell-type-specific cytokine signature genes LFC independently computed on microarray, but that unfortunately did not contain significance assessments. The considered cell-types are dendritic cells, NK, neutrophils, gd T, B, CD4T, and CD8T cells.

Starting with between-cluster analysis, we divided the cells into types using annotations from [51]. We started by applying the algorithms on a negative control setting, obtained by evenly splitting dendritic cells as two clusters (203 cells each) and comparing the two clusters (Figure 5B). As in the previous experiment, the LFC values estimated by scVI-lvm and scPhere-lvm in this negative control are significantly lower than the rest of the algorithms. We next proceeded to a comparison between different cell types (irrespective of disease status). Consistent with our results above we find that the differential expression analysis scVI-lvm and scPhere-lvm more closely match the bulk reference than the GLM-based methods, in terms of both effect sizes (LFC) and significance (ordering by FDR or p-values) estimates (Figure 5C and Figure S4 for the individual plots). We also note that the lvm based methods scale better in terms of computation time for this large dataset (Appendix J).

We next moved to the more challenging task of differential expression between the patient and control population within each cell-type. SARS-CoV-2 infection has been demonstrated to lead to aberrant secretion of pro-inflammatory cytokines such as interferon gamma [53]. As a reference to evaluate our DE analyses, we therefore used bulk transcriptome measurement of responses to different cytokines by different immune cell types. We first used a Microarray dataset [54], which provides reference LFC for cell-type-specific effects of interferon alpha. We find that scVI-lvm and scPhere-lvm better correlate with the reference microarray (significant under a t-test at significance level *α* = 0.05; Figure 5D and Figure S5 for the individual plots). We repeated this analysis with another reference dataset of bulk RNA-sequencing measurment of responses to other cytokines (IL2, IL4, IL7, IL9, IL15, and IL21), and again find that the LFC and significance evaluations by lvm-DE had consistently high correlations with the reference (Figure S6). In summary, we find that also in this more challenging task of within-cell type comparisons, lvm-DE provides more accurate estimations of gene expression compared to our benchmark methods.

## 5 Discussion

While much effort has been made recently to design deep generative models for scRNA-seq and more generally multiomics data, and despite their promises, little research has been conducted to leverage these models for DE. In this work, we present lvm-DE, a novel empirical Bayes method for differential expression, taking full advantage of the flexibility and scalability of latent variable models. This flexibility permits non-linear batch effects assumptions in the model, while global linear models for batch effect correction assume that each batch has the same cell-type proportions, often invalid in scRNA-seq experiments [8]. Furthermore, most DE pipelines, with few exceptions [55], do not take advantage of all available samples for between-type comparisons. Latent variable models that lvm-DE rely on can harness the learned shared gene-gene correlations refine gene expression modelling and hence, increase the number of detected DE genes. As such, lvm-DE helps characterize gene expression differences between cell populations in complex large-scale, batch-confounded datasets, and provides calibrated DE gene sets relying on posterior expectations of the FDP. However, our claim is not that lvm-DE can be applied to any latent variable models. In particular, we empirically observed that the library size modeling was key, e.g., in DE comparisons involving two populations with different RNA content. In scVI, the library size is a latent variable, while in ScPhere it is deterministically assigned to the number of transcripts. We noticed that latter scheme does not always successfully normalize counts, and outlined what specific elements were required to unlock differential expression for scPhere (Supplement B)

As ground-truth cell types labels are not available in most scRNA experiments, the success of DE heavily relies on a careful annotation strategy. Practical realities, including limited data availability or mislabeling, may deteriorate cell-type prediction accuracy and desired resolution, hence the range of DE analyses. The emergence of high-quality, expert-annotated cell atlases [56, 57], along with scalable transfer-learning procedures [58] show promise to provide straightforward and steady ways to annotate cells. As DE pipelines often follow the canonical approach of using the same gene counts to annotate (i.e., formulate hypotheses) and compare cell populations (testing these hypotheses), a core challenge is that the detected DE genes may reflect spurious differences, initially captured during the annotation process [59]. Fortunately, emerging approaches may limit the circularity that is central to DE for scRNA [60]. We evaluated how such a procedure, molecular cross-validation, could reduce spuriousness using synthetic data, showing marginal improvement over the canonical approach (Figure S7).

As lvm-DE outlines a generic procedure to conduct DE for latent variable models, improving the LVM of choice can be a direction to improve the quality of the predictions. One idea could be to leverage available information on gene localization to improve the underlying representation of gene-gene correlations. For instance, hmmSeq [44] is a hierarchical Bayesian model for RNA-seq taking gene positioning into account to better capture co-expression of genes in the same chromosome. Designing models that handle the sparsity of scRNA-seq experiments is another way to improve DE for latent variable models. From a biological perspective, understanding the nature of zeros and the necessity to model their excess is an active topic of research [2, 30, 31]. This question depends on the technology used for sequencing and has already been illustrated through the analysis of zero inflation for scRNA in latent variable models [15]. Besides, our aim is to formulate a general framework for differential expression for such models, that is agnostic to such modeling choices. Still, we found that lvm-DE manages to provide meaningful and calibrated decisions for zero-inflated models (Appendix I, Figure S8).

Designing better optimization procedures for latent variable models in sparse, high dimensional setups is another objective for scRNA-seq data. Sparsity can indeed hinder lvms’ ability to handle larger number of genes. In scRNA-seq data, this limitation could be overcome in several ways for models resorting on amortized variational inference. The use of more refined training procedures, relying on tighter bounds to improve the fitness of the generative model [61] could unlock DE on larger datasets. The introduction of group-sparsity priors in the decoder could effectively leverage the fact that many genes share generalized biological functions to speed up inference [62]. We hypothesize that scaling up latent variables models to larger datasets might not only help detect more DE genes, but could increase the resolution of cell populations studies.

Another extension of our work is the formulation of differential analyses for latent variable models that integrate several omic modalities. scMET [40] is a Bayesian model for differential methylation. Leveraging existing correlations [63] between methylation and gene transcription in a unified latent variable model and formulating meaningful hypotheses could be a step forward to better understand complex differentiation scenarios, e.g., for embryonic stem cells characterization [64].

## Code availability

The implementation to reproduce the experiments of this paper is available at https://github.com/PierreBoyeau/lvm-DE-reproducibility. The reference implementation of scVI is available at https://github.com/scverse/scvi-tools/tree/pierre/DE.

## Supplement

### A Implementation of the different LVMs

ScPhere was reimplemented in PyTorch using the same architecture than the original structure. However, to avoid the scalability and numerical instability of the Fisher Von-Mises distribution for consequent latent dimensions, this distribution is replaced by the Power Spherical, which does not suffer from those effects [65]. The main difference we introduced lies in the counts normalization procedure. scPhere uses observed library sizes *l*_*n*_ = Σ_*g*_ *x*_*ng*_. To ensure that the *h* are properly normalized for DE applications, we introduce a latent variable *l* accounting for library-size differences to augment scPhere. Next, we also empirically observed that the cell-specific overdispersion used by scPhere sometime caused instabilities in LFC estimation (see Appendix B), and for this reason opted for gene-specific overdispersion parameters. Originally, both scPhere and scVI use batch normalization for the encoder and decoder. After facing several stability issues in differential expression tasks with this normalization scheme, batch normalization was replaced by layer normalization. The two models originally use dropout as a regularization scheme. This choice implies an infinite ensemble of variational distributions is used during training. During evaluation on such networks, a common practice consists in disabling dropout by reweighting the regularized layers’ parameters. It is unclear why the associated distribution should be used for importance sampling. Consequently, we discard dropout layers from the architecture of the models. The remaining of the architectures follow the different models’ original implementations. We train each model with minibatches of size 1024 using the Adam optimizer to speed up convergence without computational toll. For better variational distribution coverage, we optimize the IWELBO with *K* = 25 particles, and resort to SNIS to estimate posterior expectations as described above. For these latent variable models, the optimization scheme, and in particular the learning rate is a vital hyperparameter. To avoid manual tunings of the learning rate in each experiment for each model, we rely on the automatic learning rate range test [66]. They provide data-driven, automatic learning rate choices for each latent variable model and experiment.

### B Ablation study

**Table S1:** Features ablation study. The reference model corresponds to scVI-lvm, trained with the IWELBO with 25 particles, and layer normalization. From left to right: FDR Mean absolute error (MAE; lower is better) Ranking score (RS; higher is better) PSIS shape (PSIS) LFC MSE of non-DE genes (NC, lower is better)

|  | FDR MAE | LFC MSE |
| --- | --- | --- |
| Ref. | 0.0197 | 0.0998 |
| Ref. - auto $\delta$ | 0.3185 | 0.7285 |
| Ref. - outliers | 0.3185 | 0.7285 |
| Ref. - pseudocounts | 0.0391 | 0.1201 |

First, we analyze the importance of each feature of our DE framework (Table S1). The automatic effect-size threshold and outlier removal procedures are key components to better approximate the False Discovery Rate. Using pseudo-counts slightly affects the gene rankings, but it also reduces the risk to attribute high fold changes to null genes. In the absence of this feature, the MSE of the LFC for non expressed genes nearly doubles. Pseudo-counts can hence be viewed as a regularization scheme to ensure that the detected genes have practical interest, on top of reducing the risk of predicting False Positive genes.

Next, we conduct ablation studies of the objective functions and network architectures of the different algorithms to have a clearer understanding of the hyperparameters’ influence (Tables S2 and S3). In particular, we see that the choice of overdispersion in scPhere does not modify the ability of the algorithm to properly control the FDR. However, cell-specific overdispersion seem to increase the LFC error.

**Table S2:** scVI ablation study. Differences in MAE with Table S1 are attributable to weight initialization randomness.

|  | FDR MAE | LFC MSE | PSIS |
| --- | --- | --- | --- |
| Ref. | 0.0504 | 0.1880 | 0.5988 |
| Ref. + ELBO | 0.0243 | 0.1306 | 0.4406 |

**Table S3:**
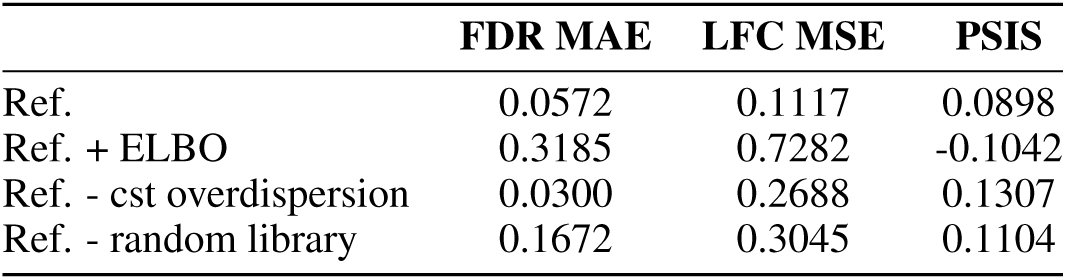
scPhere ablation study

|  | FDR MAE | LFC MSE | PSIS |
| --- | --- | --- | --- |
| Ref. | 0.0572 | 0.1117 | 0.0898 |
| Ref. + ELBO | 0.3185 | 0.7282 | -0.1042 |
| Ref. - cst overdispersion | 0.0300 | 0.2688 | 0.1307 |
| Ref. - random library | 0.1672 | 0.3045 | 0.1104 |

### C Using pseudo-counts

Most generative models rely on softmax operations to ensure that the normalized means. In such cases, the underlying expression levels have a straightforward interpretation, as they quantify gene frequencies. The softmax can however create artifacts when the two conditions *A* and *B* do not share the same library size distributions, which often is the case when *A* and *B* are cell-types. As the obtained frequencies can be arbitrarily small, the LFC distribution can reach extreme values when the gene is not expressed in both conditions.

As a solution, the offset *ϵ* (Equation 8) can serve a threshold frequency of interest that mitigates numerical artifacts. It also filters out low levels of expression, further ensuring that tagged DE genes have a practical interest. It is difficult to set the offset manually. A meaningful value depends on various parameters, including the sequencing depths of A and B, as well as the sparsity degree and shape of the data. We automatically tune this parameter, by looking at the distributions of gene expression levels for unexpressed genes in *C* ∈ {*A, B*}. Let 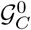 be the set of genes with zero counts in *C*, and 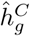 the maximum expression levels in gene *g*. If 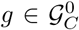, we can expect 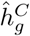 to correspond to artifacts coming from the softmax operation. Yet, it is possible that some genes were expressed, but were not observed in the counts due to the low sensitivity inherent to scRNA sequencing. To ensure the robustness of the procedure to such events, we estimate the threshold offset as the empirical *α* lower quantile of 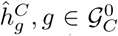, denoted as 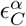. In the experiments, we use *α* = 0.90. In addition to the pseudo-count procedure, we design a robust LFC estimator that will mitigate instability when both populations do not express the gene.

### D Automatic effect-size thresholding

If is not always straightforward how *δ* should be set. The order of magnitude of the critical effect-size value that will determine if a gene is called differentially expressed depends on the two compared populations. For this reason, we suggest an automatic procedure to automatically set the value *δ*. To do so, we compute the posterior LFC medians *v*^*g*^ := median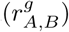) for each gene. We assume that a significant amount of LFCs will concentrate around 0 (which correspond to the genes that are equally expressed), while DE genes will concentrate around other modes. Based on this assumption, we fit a three-components Gaussian Mixture Model (GMM) based on medians *v*^*g*^. We denote *m*^*i*^, *i* ∈ {1, 2, 3} the three obtained modes. Our *δ* candidate corresponds to the mean of the largest modes (in absolute value), whose associated distributions should contain differentially expressed genes.

### E Methods detail

We provide a scaled count matrix as input (following [67]) to MAST and edgeR. DESeq2 uses the original count matrix, which is then subject to internal normalization, using the *poscounts* estimation, which seemed to improve the normalization quality. We included the batch information as covariates of the linear models included in the design matrices. Obtained p-values are corrected for multiple hypotheses testing using the Benjamini-Hochberg procedure [68] for evaluation of false discovery rate. Similar to our approach, DESeq2 formulates composite null hypotheses in which the LFC absolute value is below a certain threshold *δ*. We set the value of the *δ* parameter of DESeq2 to 0.5. To distinguish DESeq2’s predictions using composite from the point null hypotheses, we precise *δ* = 0 in the latter scenario. For other frequentist methods, which did not provide composite null hypotheses options, a subsequent filtering step of DE gene sets consisted in discarding genes with predicted LFC below 0.5 in absolute value.

### F SymSim extension to differential analysis

SymSim provides counts of the form *x*_*ng*_ along with labels *s*_*ng*_. We focus on the comparison of cell types *A* and *B*. We now describe SymSim-DA, a slight modification of the employed SymSim dataset that induces subtle differences in expression. Indeed, gene expression within a cluster can be multimodal. Better understanding how differential analysis tools handle such specificity is vital. The procedure we use consists in modifying the data to introduce multi-modalities in count distributions, from the raw counts. Let 𝒢_*d*_ be the set of differentially expressed genes between *A* and *B*. We randomly split this set in gene sets, that we denote 𝒢_*DE*_, 𝒢_*EE*_, 𝒢_*DP*_, 𝒢_*DM*_, and 𝒢_*DB*_. The first set coincides to differential expression, and hence does not require any modification of the original data. The other sets correspond to equal expression (EE), differences in proportions (DP), differences in mixture (DM), and differences in both (DB) (Figure S1A). Each set relies on a pool of held-out counts for the two cell-types to artificially introduce other modes.

Both scVI-lvm and scPhere-lvm compare favorably to the other algorithms (Figure S1B.).

**Figure S1:**
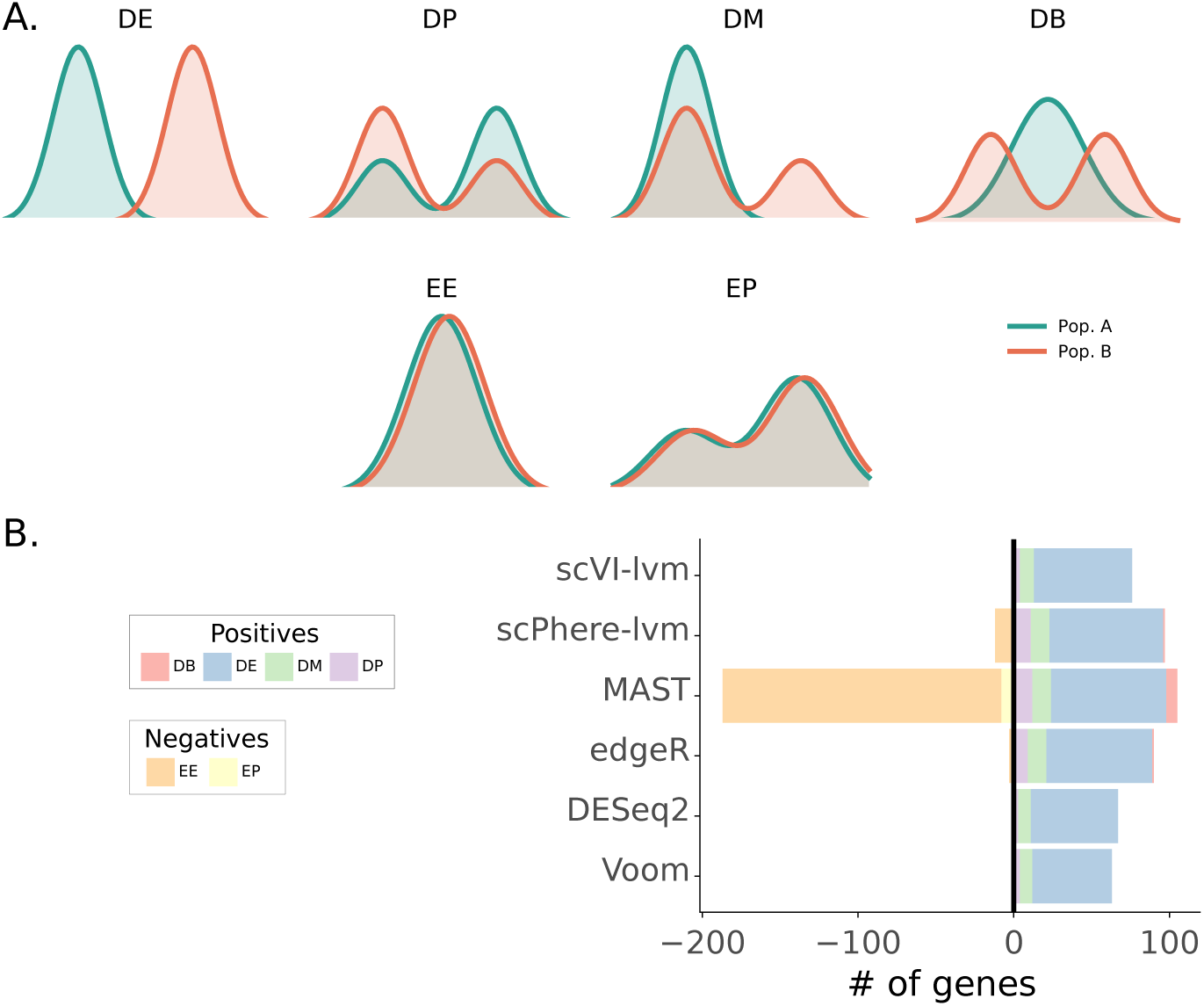
SymSim-DA results, extending the dataset to unlock a broader differential analysis scenario. **A**. top: Overview of the different differential distributions for differential analysis. From left to right: 1. Differential expression, in which the modes of gene expressions are different. 2. Differences in proportion, in which both populations express the gene according to two modes, but with different weights 3. Differences in mixture, in which the two populations share the same mode but one of them also has another mode 4. Differences in both, a combination of the last two scenarios. bottom: differential distributions that do not induce differences between the populations. In both cases, both populations have the same gene expression, but in EP the gene expression is multimodal. **B**. Types of genes detected by each method (at FDR level 0.05). Detections corresponding to DB, DE, DM, or DP correspond to true positives, while EE or EP genes denote false positives.

### G Differential expression on MUSCAT

We proceed to the same analysis than the one conducted in SymSim for a synthetic dataset generated using MUSCAT (Figure S2). In this framework, an external dataset helps to fit sample-specific negative binomials (with shared, gene specific overdispersion parameters). These reference distributions are used to construct a synthetic dataset consisting of several subpopulations. A first simple scenario consists in only focusing on differences of expression. In this case, each subpopulation will follow negative binomial count distributions, with subpopulation-specific mean parameters. Here, the presence of other cell-types cannot be leveraged to improve differential expression performance between A and B S2C). While lvm-DE’s FDR control is acceptable, linear models also work well in this scenario (Figure S2D).

### H Individual scatter plots

We here show the complete inter cell-types analysis for the PBMC (Figure S3) and Blish (Figure S4) datasets, as well as the intra cell-types analysis (Figure S5).

### I Zero-inflation effects

We here discuss to the flexibility of the lvm-DE framework to changes in the noise model. In particular, we investigate if zero-inflated models still control the FDR in the lvm-DE framework (Figure S8). Originally designed to better account for the low sensitivity of droplet-based sequencing procedures, its pertinence to properly model scRNA has been challenged [31]. In this dataset, we find that a zero-inflated negative binomial likelihood does not seem to better capture differentially expressed genes that their negative binomial counterparts. However, zero-inflated models may still be used by our model.

**Figure S2:**
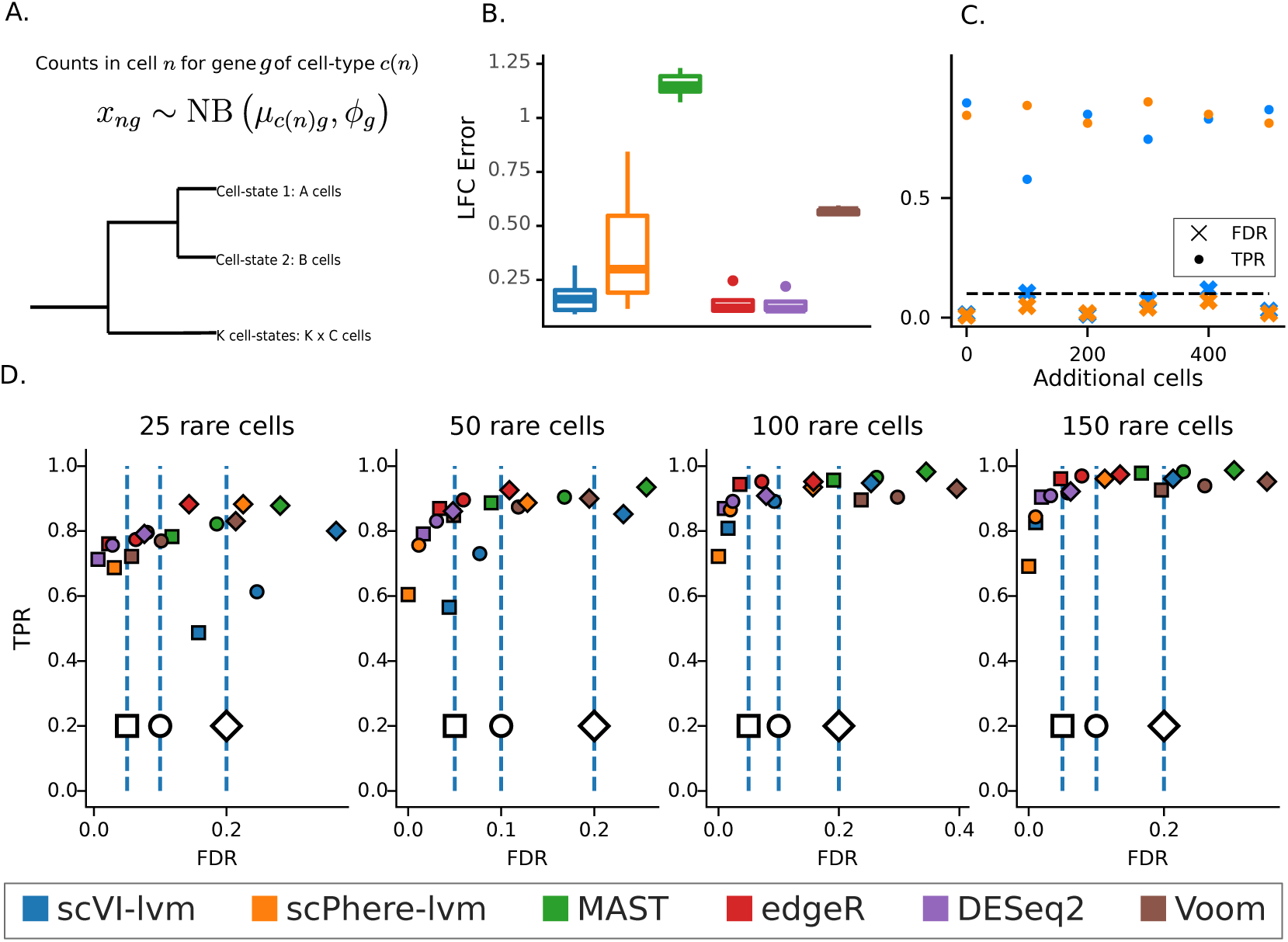
MUSCAT results. **A**. Dataset presentation. **B**. LFC point estimation error when comparing two populations of 200 cells. For Bayesian techniques we summarize the posterior LFC distribution by its median. **C**. TPR (*dots*) and FDR (*crosses*) changes for an increasing number of external cells for the different latent variable models. **D**. FDR and TPR of decisions for the detection of DE genes when comparing 25, 50, 100, and 150 cells to 500 cells. Squares, circles and losanges correspond to decision controlling FDR at targets 0.05, 0.1, and 0.2. Because the original scVI model does not provide explicit thresholds to tag DE genes, we reject the null when Bayes factors are greater that 3 in absolute value.

### J Running times

Running times of the different methods are provided in Table S4. They correspond for the inter cell-type comparisons performed on the Blish datasets. All experiments were run on a remote server with one Tesla V100 GPU, 40 virtual processors and 120 GB of RAM. We observed that the GLMs scaled poorly with higher numbers of batches, with the exception of Voom, which however provided poor correlation with bulk-RNA. While both scVI and scVI-DE require additional time to fit their generative model on the entire dataset, the lvm framework also provides other functionalities (clustering, normalization, imputation tasks) often required for differential expression analysis that require the use of external packages for the other DE protocols. In addition, scVI-DE differential expression execution time is constant with respect to the number of cells analyzed for differential expression.

**Table S4:** Mean Running time on the Blish dataset for the comparison of the six pairs of cell types. Training times correspond to the required time to fit latent variable models on the whole dataset for 250 epochs.

| Blish | DESeq2 | MAST | edgeR | Voom | scVI-lvm | scPhere-lvm |
| --- | --- | --- | --- | --- | --- | --- |
| Pre-training (Minutes) | N/A | N/A | N/A | N/A | 21.4 | 23.4 |
| Differential expression (Seconds) | 266.7 | 139.0 | 102.7 | 1105.3 | 35.1 | 41.4 |

**Figure S3:**
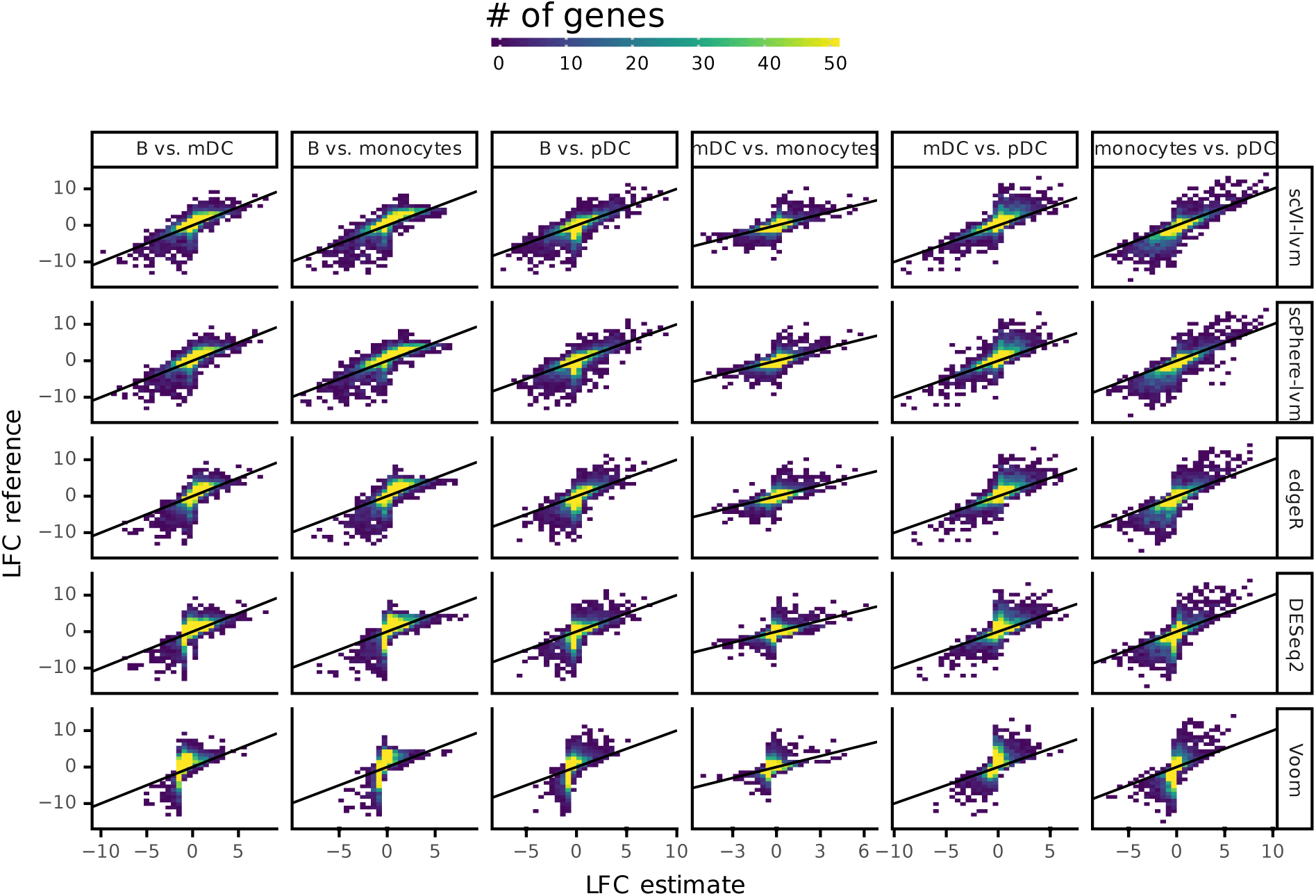
Inter cell-types analysis for the PBMC dataset, through the comparison of predicted LFC with reference bulk for pairwise comparisons of B cells, mDC, pDC and monocytes.

**Figure S4:**
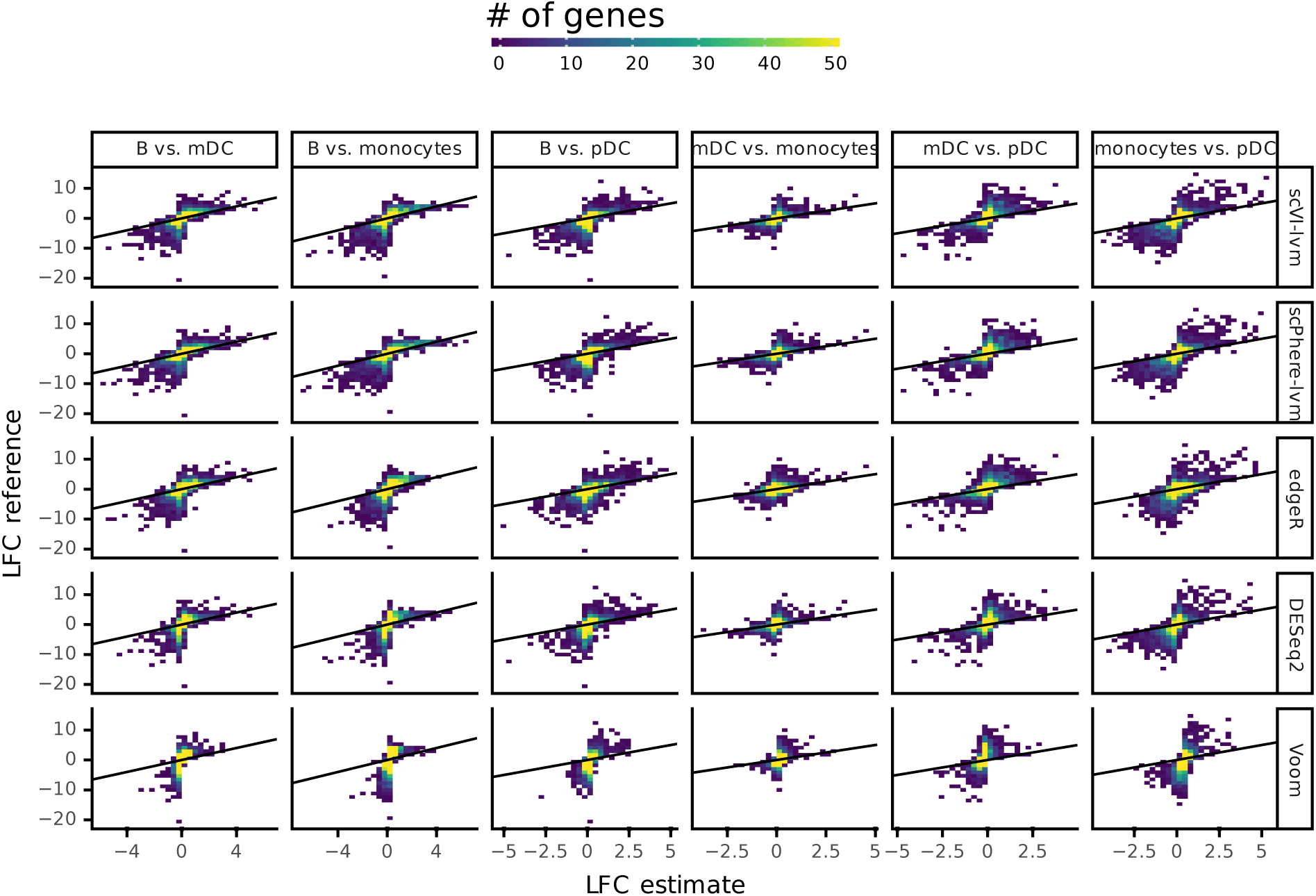
Inter cell-types analysis for the Blish dataset, through the comparison of predicted LFC with reference bulk for pairwise comparisons of B cells, mDC, pDC and monocytes.

**Figure S5:**
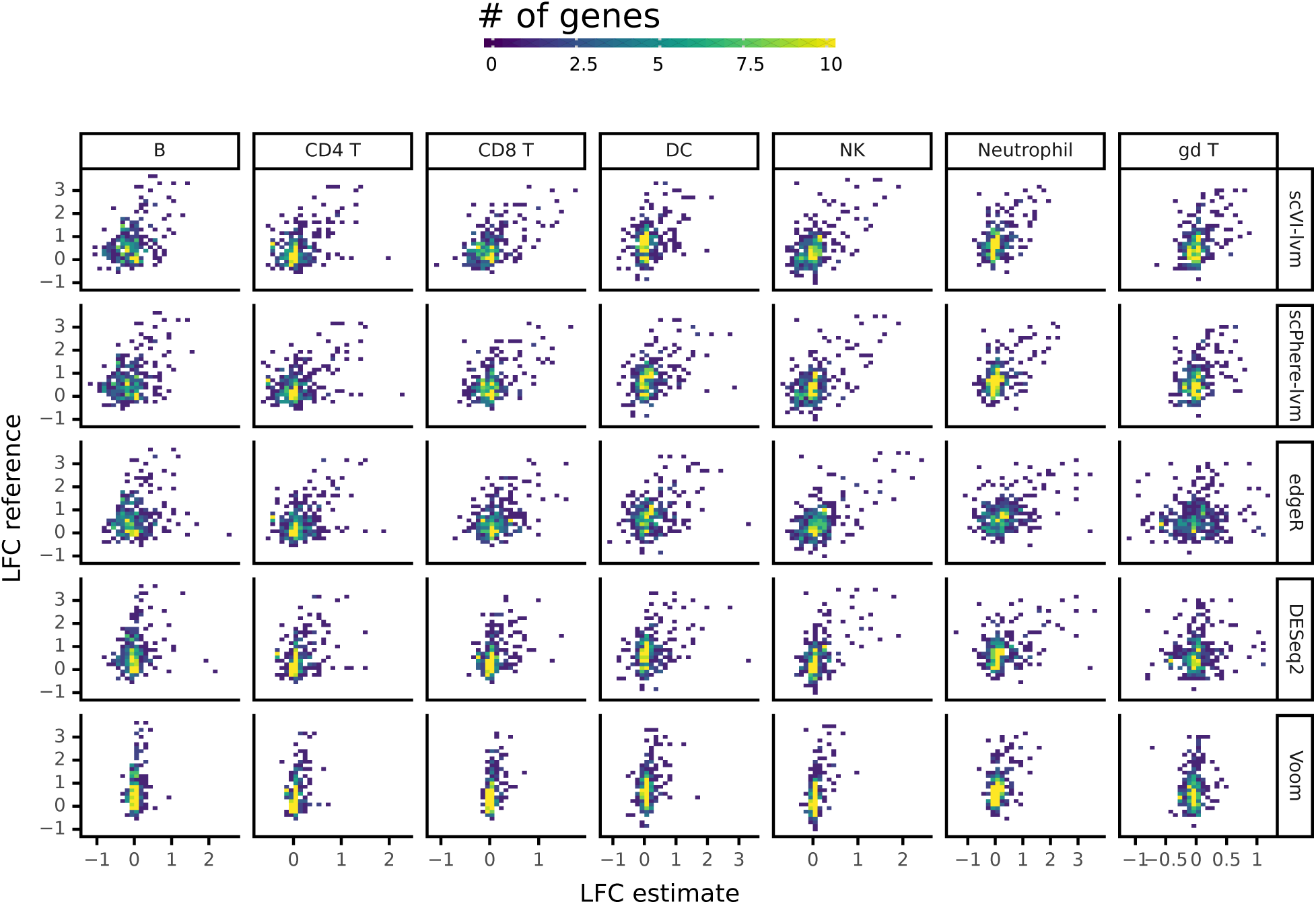
Within cell-types analysis for the Blish dataset, through the comparison of predicted LFC with cytokyne genes microarrray.

**Figure S6:**
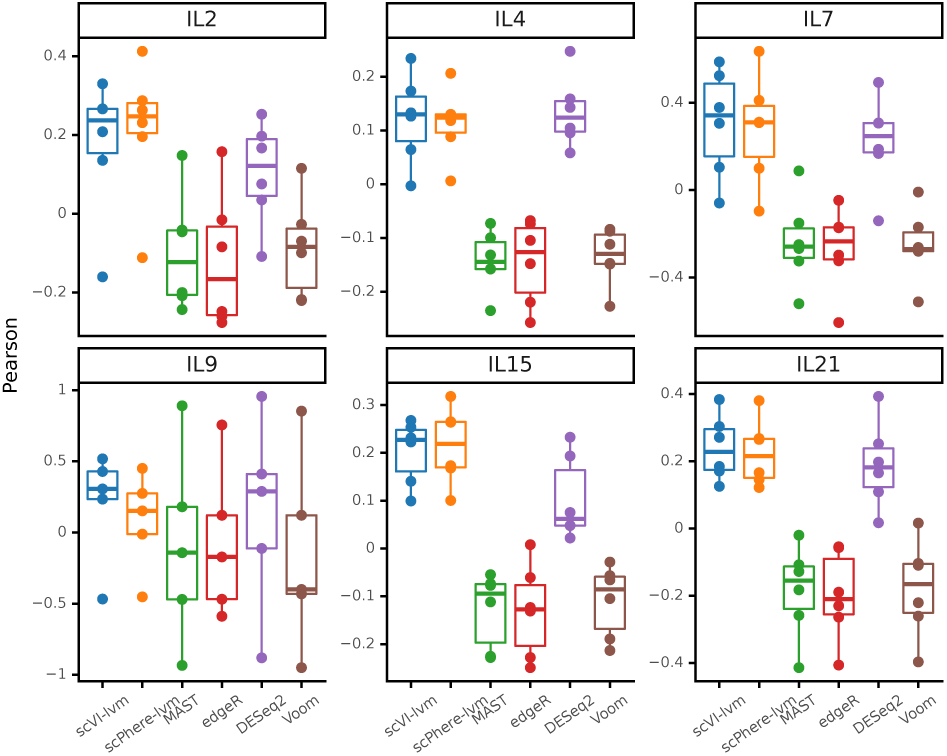
Predicted LFC correlations with reference RNA data for several cell-type-specific cytokine stimulation markers. In each subgraph, points correspond to the correlation for a given cell-type (B, Dendritic, NK, CD4, CD8, or gamma-deltas.)

**Figure S7:**
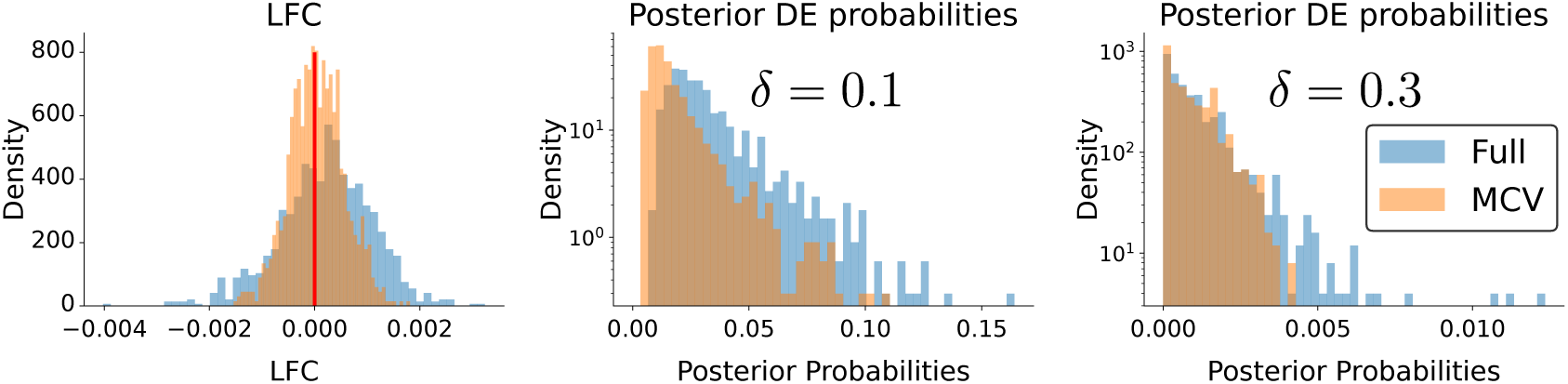
Effect of molecular cross-validation (MCV) on LFC estimation and posterior DE probabilities on Poisson negative-control data using scVI-lvm. The model either used the same transcripts to train the model and to detect DE genes (Full) or used different molecules for model fitting and inference (MCV), according to a 80%-20% split. *Left*: Distribution of obtained LFC estimates. *Middle* and *right*: Distribution of posterior DE probabilities for two LFC thresholds (*δ* = 0.1 and *δ* = 0.3).

**Figure S8:**
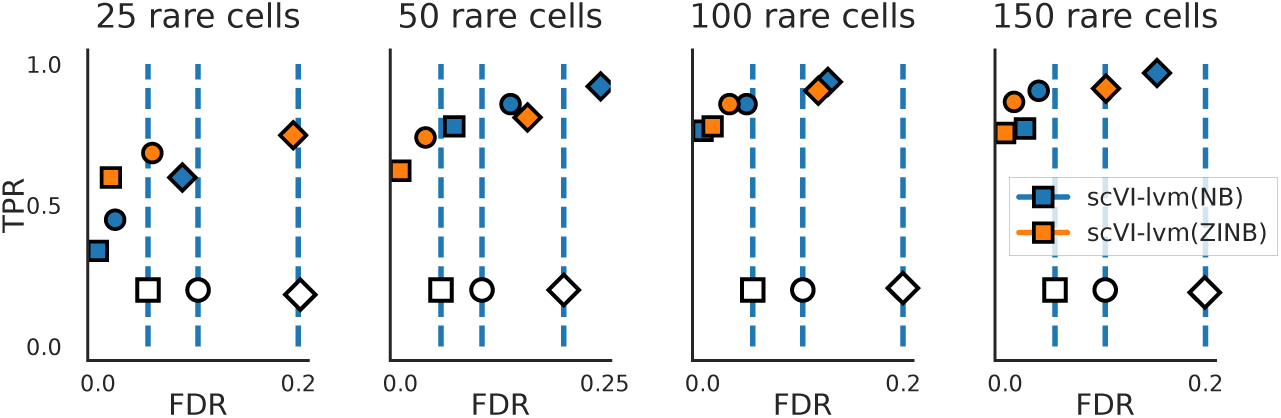
FDR-TPR curves on the Symsim dataset for varying *A* sizes, comparing different likelihood models for scVI-lvm. In this experiment, *B* = 500, *C* = 2000.

## References

[1] Allon Wagner, Aviv Regev, and Nir Yosef. Revealing the vectors of cellular identity with single-cell genomics. Nature biotechnology, 34(11):1145–1160, 2016.

[2] Greg Finak, Andrew McDavid, Masanao Yajima, Jingyuan Deng, Vivian Gersuk, Alex K Shalek, Chloe K Slichter, Hannah W Miller, M Juliana McElrath, Martin Prlic, et al. MAST: a flexible statistical framework for assessing transcriptional changes and characterizing heterogeneity in single-cell RNA sequencing data. Genome biology, 16(1):1–13, 2015.

[3] Peter V Kharchenko, Lev Silberstein, and David T Scadden. Bayesian approach to single-cell differential expression analysis. Nature methods, 11(7):740–742, 2014.

[4] Rahul Satija, Jeffrey A Farrell, David Gennert, Alexander F Schier, and Aviv Regev. Spatial reconstruction of single-cell gene expression data. Nature biotechnology, 33(5):495–502, 2015.

[5] Michael I Love, Wolfgang Huber, and Simon Anders. Moderated estimation of fold change and dispersion for RNA-seq data with DESeq2. Genome biology, 15(12):1–21, 2014.

[6] Mark D Robinson, Davis J McCarthy, and Gordon K Smyth. edgeR: a bioconductor package for differential expression analysis of digital gene expression data. Bioinformatics, 26(1):139–140, 2010.

[7] Jordan W Squair, Matthieu Gautier, Claudi Kathe, Mark A Anderson, Nicholas D James, Thomas H Hutson, Rémi Hudelle, Taha Qaiser, Kaya JE Matson, Quentin Barraud, et al. Confronting false discoveries in single-cell differential expression. bioRxiv, 2021.

[8] Laleh Haghverdi, Aaron TL Lun, Michael D Morgan, and John C Marioni. Batch effects in single-cell RNA-sequencing data are corrected by matching mutual nearest neighbors. Nature biotechnology, 36(5):421–427, 2018.

[9] Malte D Luecken, Maren Buttner, Kridsadakorn Chaichoompu, Anna Danese, Marta Interlandi, Michaela F Müller, Daniel C Strobl, Luke Zappia, Martin Dugas, Maria Colomé-Tatché, et al. Benchmarking atlas-level data integration in single-cell genomics. BioRxiv, 2020.

[10] W. Evan Johnson, Cheng Li, and Ariel Rabinovic. Adjusting batch effects in microarray expression data using empirical Bayes methods. Biostatistics, 8(1):118–127, 04 2006.

[11] Diederik P Kingma and Max Welling. Auto-encoding variational bayes. arXiv preprint 1312.6114, 2013.

[12] Romain Lopez, Adam Gayoso, and Nir Yosef. Enhancing scientific discoveries in molecular biology with deep generative models. Molecular Systems Biology, 16(9):e9198, 2020.

[13] Jiarui Ding and Aviv Regev. Deep generative model embedding of single-cell RNA-seq profiles on hyperspheres and hyperbolic spaces. Nature Communications, 2021.

[14] Romain Lopez, Jeffrey Regier, Michael B Cole, Michael I Jordan, and Nir Yosef. Deep generative modeling for single-cell transcriptomics. Nature methods, 15(12):1053–1058, 2018.

[15] Chenling Xu, Romain Lopez, Edouard Mehlman, Jeffrey Regier, Michael I Jordan, and Nir Yosef. Probabilistic harmonization and annotation of single-cell transcriptomics data with deep generative models. Molecular systems biology, 17(1):e9620, 2021.

[16] Mohammad Lotfollahi, F Alexander Wolf, and Fabian J Theis. scGen predicts single-cell perturbation responses. Nature methods, 16(8):715–721, 2019.

[17] Stephen J Fleming, John C Marioni, and Mehrtash Babadi. Cellbender remove-background: a deep generative model for unsupervised removal of background noise from scRNA-seq datasets. BioRxiv, page 791699, 2019.

[18] Adam Gayoso, Romain Lopez, Galen Xing, Pierre Boyeau, Valeh Valiollah Pour Amiri, Justin Hong, Katherine Wu, Michael Jayasuriya, Edouard Mehlman, Maxime Langevin, Yining Liu, Jules Samaran, Gabriel Misrachi, Achille Nazaret, Oscar Clivio, Chenling Xu, Tal Ashuach, Mariano Gabitto, Mohammad Lotfollahi, Valentine Svensson, Eduardo da Veiga Beltrame, Vitalii Kleshchevnikov, Carlos Talavera-López, Lior Pachter, Fabian J. Theis, Aaron Streets, Michael I. Jordan, Jeffrey Regier, and Nir Yosef. A python library for probabilistic analysis of single-cell omics data. Nature Biotechnology, Feb 2022.

[19] Dominic Grün, Lennart Kester, and Alexander Van Oudenaarden. Validation of noise models for single-cell transcriptomics. Nature methods, 11(6):637–640, 2014.

[20] Charlotte Soneson and Mark D Robinson. Bias, robustness and scalability in single-cell differential expression analysis. Nature methods, 15(4):255, 2018.

[21] Charity W Law, Yunshun Chen, Wei Shi, and Gordon K Smyth. voom: Precision weights unlock linear model analysis tools for RNA-seq read counts. Genome biology, 15(2):1–17, 2014.

[22] Peng Qiu. Embracing the dropouts in single-cell rna-seq analysis. Nature communications, 11(1):1–9, 2020.

[23] Valentine Svensson, Eduardo da Veiga Beltrame, and Lior Pachter. A curated database reveals trends in single-cell transcriptomics. Database, 2020.

[24] Sabrina Rashid, Sohrab Shah, Ziv Bar-Joseph, and Ravi Pandya. Dhaka: variational autoencoder for unmasking tumor heterogeneity from single cell genomic data. Bioinformatics, 2019.

[25] Dongfang Wang and Jin Gu. VASC: dimension reduction and visualization of single-cell RNA-seq data by deep variational autoencoder. Genomics, proteomics & bioinformatics, 16(5):320–331, 2018.

[26] Gökcen Eraslan, Lukas M Simon, Maria Mircea, Nikola S Mueller, and Fabian J Theis. Single-cell RNA-seq denoising using a deep count autoencoder. Nature communications, 10(1):1–14, 2019.

[27] Christopher Heje Grønbech, Maximillian Fornitz Vording, Pascal N Timshel, Casper Kaae Sønderby, Tune H Pers, and Ole Winther. scVAE: Variational auto-encoders for single-cell gene expression data. Bioinformatics, 36(16):4415–4422, 2020.

[28] Matthew Amodio, David Van Dijk, Krishnan Srinivasan, William S Chen, Hussein Mohsen, Kevin R Moon, Allison Campbell, Yujiao Zhao, Xiaomei Wang, Manjunatha Venkataswamy, et al. Exploring single-cell data with deep multitasking neural networks. Nature methods, 16(11):1139–1145, 2019.

[29] Mohammad Lotfollahi, Mohsen Naghipourfar, Fabian J Theis, and F Alexander Wolf. Conditional out-of-sample generation for unpaired data using trVAE. arXiv preprint 1910.01791, 2019.

[30] Oscar Clivio, Romain Lopez, Jeffrey Regier, Adam Gayoso, Michael I Jordan, and Nir Yosef. Detecting zero-inflated genes in single-cell transcriptomics data. bioRxiv, page 794875, 2019.

[31] Valentine Svensson. Droplet scRNA-seq is not zero-inflated. Nature Biotechnology, 38(2):147–150, 2020.

[32] Tallulah S Andrews and Martin Hemberg. False signals induced by single-cell imputation. F1000Research, 7, 2018.

[33] Valentine Svensson, Adam Gayoso, Nir Yosef, and Lior Pachter. Interpretable factor models of single-cell rna-seq via variational autoencoders. Bioinformatics, 36(11):3418–3421, 2020.

[34] Erik Nijkamp, Bo Pang, Tian Han, Linqi Zhou, Song-Chun Zhu, and Ying Nian Wu. Learning multi-layer latent variable model via variational optimization of short run mcmc for approximate inference. In European Conference on Computer Vision, pages 361–378. Springer, 2020.

[35] Romain Lopez, Pierre Boyeau, Nir Yosef, Michael I Jordan, and Jeffrey Regier. Decision-making with autoencoding variational Bayes. Advances in Neural Information Processing Systems, 2020.

[36] Justin Domke and Daniel R Sheldon. Importance weighting and variational inference. In S. Bengio, H. Wallach, H. Larochelle, K. Grauman, N. Cesa-Bianchi, and R. Garnett, editors, Advances in Neural Information Processing Systems, volume 31. Curran Associates, Inc., 2018.

[37] Yuling Yao, Aki Vehtari, Daniel Simpson, and Andrew Gelman. Yes, but did it work?: Evaluating variational inference. In International Conference on Machine Learning, pages 5581–5590. PMLR, 2018.

[38] Jianxing Feng, Clifford A Meyer, Qian Wang, Jun S Liu, X Shirley Liu, and Yong Zhang. GFOLD: a generalized fold change for ranking differentially expressed genes from RNA-seq data. Bioinformatics, 28(21):2782–2788, 2012.

[39] Malte D Luecken and Fabian J Theis. Current best practices in single-cell RNA-seq analysis: a tutorial. Molecular systems biology, 15(6):e8746, 2019.

[40] Chantriolnt-Andreas Kapourani, Ricard Argelaguet, Guido Sanguinetti, and Catalina A Vallejos. scMET: Bayesian modelling of DNA methylation heterogeneity at single-cell resolution. bioRxiv, 2020.

[41] James O Berger. Statistical decision theory and Bayesian analysis. Springer Science & Business Media, 2013.

[42] Michael A Newton, Amine Noueiry, Deepayan Sarkar, and Paul Ahlquist. Detecting differential gene expression with a semiparametric hierarchical mixture method. Biostatistics, 5(2):155–176, 2004.

[43] Juhee Lee, Yuan Ji, Shoudan Liang, Guoshuai Cai, and Peter Müller. On differential gene expression using RNA-seq data. Cancer informatics, 10:CIN–S7473, 2011.

[44] Shiqi Cui, Subharup Guha, Marco AR Ferreira, Allison N Tegge, et al. hmmSeq: A hidden Markov model for detecting differentially expressed genes from RNA-seq data. The Annals of Applied Statistics, 9(2):901–925, 2015.

[45] Xiuwei Zhang, Chenling Xu, and Nir Yosef. Simulating multiple faceted variability in single cell RNA sequencing. Nature communications, 10(1):1–16, 2019.

[46] Keegan D Korthauer, Li-Fang Chu, Michael A Newton, Yuan Li, James Thomson, Ron Stewart, and Christina Kendziorski. A statistical approach for identifying differential distributions in single-cell rna-seq experiments. Genome biology, 17(1):1–15, 2016.

[47] Helena L Crowell, Charlotte Soneson, Pierre-Luc Germain, Daniela Calini, Ludovic Collin, Catarina Raposo, Dheeraj Malhotra, and Mark D Robinson. muscat detects subpopulation-specific state transitions from multisample multi-condition single-cell transcriptomics data. Nature communications, 11(1):1–12, 2020.

[48] Grace XY Zheng, Jessica M Terry, Phillip Belgrader, Paul Ryvkin, Zachary W Bent, Ryan Wilson, Solongo B Ziraldo, Tobias D Wheeler, Geoff P McDermott, Junjie Zhu, et al. Massively parallel digital transcriptional profiling of single cells. Nature communications, 8(1):1–12, 2017.

[49] Gianni Monaco, Bernett Lee, Weili Xu, Seri Mustafah, You Yi Hwang, Christophe Carre, Nicolas Burdin, Lucian Visan, Michele Ceccarelli, Michael Poidinger, et al. RNA-seq signatures normalized by mRNA abundance allow absolute deconvolution of human immune cell types. Cell reports, 26(6):1627–1640, 2019.

[50] Tamim Abdelaal, Lieke Michielsen, Davy Cats, Dylan Hoogduin, Hailiang Mei, Marcel JT Reinders, and Ahmed Mahfouz. A comparison of automatic cell identification methods for single-cell RNA sequencing data. Genome biology, 20(1):1–19, 2019.

[51] Aaron J Wilk, Arjun Rustagi, Nancy Q Zhao, Jonasel Roque, Giovanny J Martínez-Colón, Julia L McKechnie, Geoffrey T Ivison, Thanmayi Ranganath, Rosemary Vergara, Taylor Hollis, et al. A single-cell atlas of the peripheral immune response in patients with severe COVID-19. Nature medicine, 26(7):1070–1076, 2020.

[52] Elior Rahmani, Michael I Jordan, and Nir Yosef. Identifying systematic variation at the single-cell level by leveraging low-resolution population-level data. bioRxiv, 2022.

[53] Antonella Fara, Zan Mitrev, Rodney Alexander Rosalia, and Bakri M Assas. Cytokine storm and covid-19: a chronicle of pro-inflammatory cytokines. Open biology, 10(9):200160, 2020.

[54] Tracy SP Heng, Michio W Painter, Kutlu Elpek, Veronika Lukacs-Kornek, Nora Mauermann, Shannon J Turley, Daphne Koller, Francis S Kim, Amy J Wagers, Natasha Asinovski, et al. The immunological genome project: networks of gene expression in immune cells. Nature immunology, 9(10):1091–1094, 2008.

[55] Vasilis Ntranos, Lynn Yi, Páll Melsted, and Lior Pachter. A discriminative learning approach to differential expression analysis for single-cell RNA-seq. Nature methods, 16(2):163–166, 2019.

[56] Stephen R Quake, Tabula Sapiens Consortium, et al. The tabula sapiens: a single cell transcriptomic atlas of multiple organs from individual human donors. Biorxiv, 2021.

[57] Herbert B Schiller, Daniel T Montoro, Lukas M Simon, Emma L Rawlins, Kerstin B Meyer, Maximilian Strunz, Felipe A Vieira Braga, Wim Timens, Gerard H Koppelman, GR Scott Budinger, et al. The human lung cell atlas: a high-resolution reference map of the human lung in health and disease. American journal of respiratory cell and molecular biology, 61(1):31–41, 2019.

[58] Mohammad Lotfollahi, Mohsen Naghipourfar, Malte D Luecken, Matin Khajavi, Maren Büttner, Marco Wagenstetter, Z. iga Avsec, Adam Gayoso, Nir Yosef, Marta Interlandi, et al. Mapping single-cell data to reference atlases by transfer learning. Nature Biotechnology, 40(1):121–130, 2022.

[59] Nikolaus Kriegeskorte, W Kyle Simmons, Patrick SF Bellgowan, and Chris I Baker. Circular analysis in systems neuroscience: the dangers of double dipping. Nature neuroscience, 12(5):535–540, 2009.

[60] Joshua Batson, Löic Royer, and James Webber. Molecular cross-validation for single-cell rna-seq. BioRxiv, page 786269, 2019.

[61] Rahul Krishnan, Dawen Liang, and Matthew Hoffman. On the challenges of learning with inference networks on sparse, high-dimensional data. In International Conference on Artificial Intelligence and Statistics, pages 143–151. PMLR, 2018.

[62] Mohammad Lotfollahi, Sergei Rybakov, Karin Hrovatin, Soroor Hediyeh-zadeh, Carlos Talavera-López, Alexander Misharin, and Fabian J Theis. Biologically informed deep learning to infer gene program activity in single cells. bioRxiv, 2022.

[63] Jeongwoo Lee, Daehee Hwang, et al. Single-cell multiomics: technologies and data analysis methods. Experimental & Molecular Medicine, 52(9):1428–1442, 2020.

[64] Stephen J Clark, Ricard Argelaguet, Chantriolnt-Andreas Kapourani, Thomas M Stubbs, Heather J Lee, Celia Alda-Catalinas, Felix Krueger, Guido Sanguinetti, Gavin Kelsey, John C Marioni, et al. scNMT-seq enables joint profiling of chromatin accessibility DNA methylation and transcription in single cells. Nature communications, 9(1):1–9, 2018.

[65] Nicola De Cao and Wilker Aziz. The power spherical distribution. arXiv preprint 2006.04437, 2020.

[66] Leslie N Smith. Cyclical learning rates for training neural networks. In 2017 IEEE winter conference on applications of computer vision (WACV), pages 464–472. IEEE, 2017.

[67] Mark D Robinson and Alicia Oshlack. A scaling normalization method for differential expression analysis of RNA-seq data. Genome biology, 11(3):1–9, 2010.

[68] Yoav Benjamini and Yosef Hochberg. Controlling the false discovery rate: a practical and powerful approach to multiple testing. Journal of the Royal statistical society: series B (Methodological), 57(1):289–300, 1995.

